# Human thymopoiesis produces polyspecific CD8^+^ α/β T cells responding to multiple viral antigens

**DOI:** 10.1101/2020.07.27.223354

**Authors:** Valentin Quiniou, Pierre Barennes, Vanessa Mhanna, Paul Stys, Hélène Vantomme, Zhicheng Zhou, Federica Martina, Nicolas Coatnoan, Michèle Barbié-Sastre, Hang-Phuong Pham, Béatrice Clemenceau, Henri Vié, Mikhail Shugay, Adrien Six, Barbara Brandao, Roberto Mallone, Encarnita Mariotti-Ferrandiz, David Klatzmann

## Abstract

T cell receptors (TCRs) are formed by stochastic gene rearrangements, theoretically generating >10^19^ sequences. They are selected during thymopoiesis, which releases a repertoire of about 10^8^ unique TCRs per individual. How evolution shaped a process that produces TCRs that can effectively handle a countless and evolving set of infectious agents is a central question of immunology. The paradigm is that a diverse enough repertoire of TCRs should always provide a proper, though rare, specificity for any given need. Expansion of such rare T cells would provide enough fighters for an effective immune response and enough antigen-experienced cells for memory. We show here that human thymopoiesis releases a large population of CD8^+^ T cells harboring α/β paired TCRs that (i) have high generation probabilities and (ii) a preferential usage of some V and J genes, (iii) are shared between individuals and (iv) can each recognize and be activated by multiple unrelated viral peptides, notably from EBV, CMV and influenza. These polyspecific T cells may represent a first line of defense that is mobilized in response to infections before a more specific response subsequently ensures viral elimination. Our results support an evolutionary selection of polyspecific α/β TCRs for broad antiviral responses and heterologous immunity.

## INTRODUCTION

Specificity is considered a hallmark of the adaptive immune response. For T cells, specificity is mediated by T cell receptors (TCRs), which interact with peptides presented by major histocompatibility complex (MHC) molecules through their complementary-determining-region-3 (CDR3). TCRs are formed by rearrangements between hundreds of gene segments at the alpha and beta loci, theoretically generating >10^60^ possible sequences (*1–3*). Each rearrangement has its probability of generation, which has been modeled to vary by about 10 orders of magnitude for the beta chain alone (*4*). During their development within the thymus, T cells undergo a selection largely based on the strength of their activation by thymic antigen-presenting cells that eliminates >80% of them (*5–7*). For each individual, this process releases a pool of T cells expressing a repertoire of approximately 10^8^ unique TCRs (*8*).

How evolution shaped a process that selects cells that would efficiently respond to antigens that are not yet present, i.e. from an infectious agent to come, is a central question of immunology. The paradigm is that the production and selection of a diverse enough repertoire of TCRs should always provide a proper, though rare, specificity for any given need. Expansion of such rare T cells would then provide enough fighters for an effective immune response and later enough antigen-experienced cells for immune memory (*9*). Thus, it is widely accepted that “*T cell-mediated immunity to infection is due to the proliferation and differentiation of rare clones in the preimmune repertoire that by chance express TCRs specific for peptide-MHC (pMHC) ligands derived from the microorganism*”(*10*).

However, this theory is challenged by overlooked data questioning the essence of specificity. First, a child with an X-linked severe combined immunodeficiency who had a reverse mutation in a single T cell progenitor was protected from infections despite having a very restricted repertoire of only about 1,000 different TCRs (*11*). How such a repertoire (10^−5^ of a normal repertoire size) could handle the numerous infections occurring during childhood awaits explanation. Second, robust experimental and epidemiological data have highlighted the importance of heterologous immunity, i.e. immunity to one pathogen afforded by the exposure to unrelated pathogens (*12–14*). Likewise, for example, (i) memory T cells that are specific for one virus can become activated during infection with an unrelated virus (*15*), (ii) there are numerous virus-specific memory phenotype T cells in unexposed individuals (*16*), (iii) CMV infection enhances the immune response to influenza (*17*) and (iv) tumor-resident memory HBV-specific T cell responses correlate with hepatocellular carcinoma outcome(18).

We hypothesized that high-resolution sequencing of the TCR repertoire of developing thymocytes and peripheral T cells would yield insights into repertoire selection and function towards pathogens. We focused on the development of CD8^+^ T cells as they represent the main effectors against viral pathogens, those that have had most chances to influence the evolution of our immune system. We obtained samples from organ donors allowing access to thymocytes and peripheral T cells, as well as datasets from patients undergoing vaccinations or with infections; we also used TCR data available from multiple public repositories.

We identified a population of CD8^+^ T cells harboring diversified and polyspecific α/β TCRs that each can bind to multiple unrelated viral peptides, notably those from commonly encountered viruses such as EBV, CMV or influenza. We hypothesize that these cells represent a first line of defense that is promptly mobilized in response to infections, possibly containing them before a more specific response subsequently ensures their control. Our results support an evolutionary selection of polyspecific α/βTCRs for broad antiviral responses and heterologous immunity.

## RESULTS

### Thymopoiesis selects a large and diverse set of clustered CDR3s with high generation probabilities

To investigate how TCR diversity influences immune response to infectious agents, we started by analyzing the TCR repertoire dynamics of developing thymocytes. We first focused on the hypervariable CDR3 region of the TCR that interacts with the antigenic peptide, while CDR1 and CDR2 usually interact with HLA molecules (*19*). CDR3 analyses can therefore be used to investigate the sharing of TCR specificities across individuals with distinct HLA molecules. We analyzed the repertoire of purified CD4^+^CD8^+^CD3^−^ (DPCD3^−^), CD4^+^CD8^+^CD3^+^ (DPCD3^+^) and CD4^−^ CD8^+^CD3^+^ (CD8^+^) thymocytes (Fig. S1A). DPCD3^−^ thymocytes represent the earliest stage of TCR β-chain gene recombination, and their repertoire embodies the unaltered outcome of the TCR generation process; DPCD3^+^ thymocytes are at an early stage of the selection process and their repertoire should be minimally modified; CD8^+^ thymocytes have passed the selection process and bear a fully selected repertoire. We analyzed and represented the structure of these repertoires by connecting CDR3s (nodes) differing by at most one single amino acid (AA) (Levenshtein distance less than or equal to one: LD≤1). LD≤1 connected CDR3s have been described to most often bind the same peptide (*20–26*) (Fig. S1B). In such networks, connected CDR3s are designated as *clustered nodes* and the others as *dispersed nodes*. For normalization, we represented the first 18,000 most expressed β or α CDR3s from each sample.

We observed a significant increase in the number of clustered βCDR3s from DPCD3^−^ to CD8^+^ thymocytes (Fig. 1A and B) (p<0.0001), which was remarkably consistent among all individuals studied, independently of their age, sex or HLA (Fig. S2). Node’s degree, i.e., its number of CDR3 neighbors based on LD≤1, was also significantly increased during T cell differentiation for clustered βCDR3s (Fig. 1C and D) (p<0.0001). Similar results were found for the αCDR3s (Fig. S3A-B). These observations suggest a positive selection during thymopoiesis of TCRs with close CDR3 sequences, and therefore with shared recognition properties.

**Figure 1.**
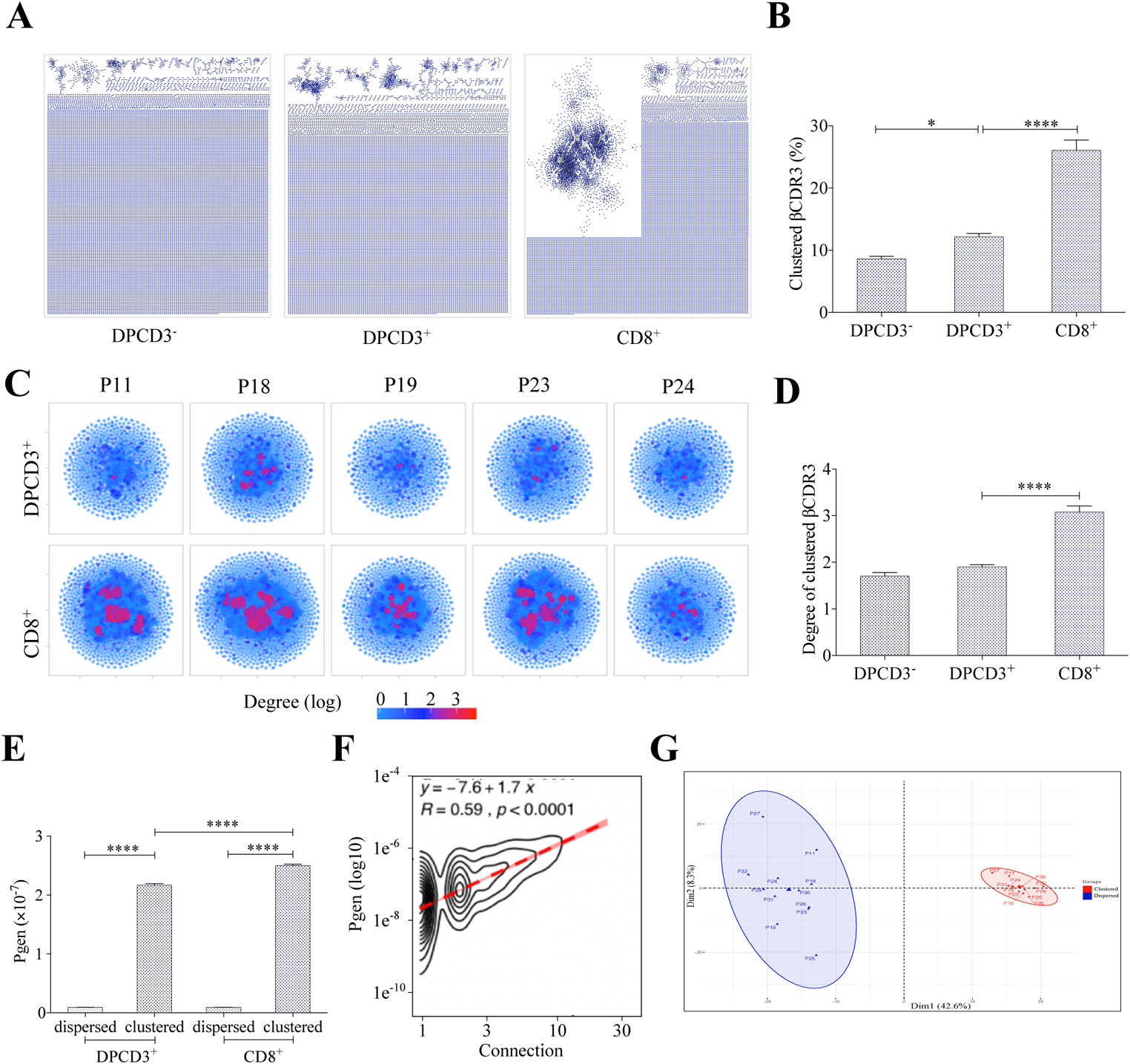
Thymocyte differentiation produces clustered CDR3s with high generation probability and preferential TRB VJ gene combinations. A. Representation of βCDR3aa networks from DPCD3^−^, DPCD3^+^ and CD8+. Each dot represents a single CDR3. Dot are connected (forming clusters) by edges defined by Levenshtein distance of ≤1 (one AA substitution/insertion/deletion).
B. Percentage of clustered βCDR3aa. (*p=0.0152 and ****p<0.0001, Mann-Whitney test).
C. βCDR3aa clustered from DPCD3^+^ and ThyCD8. Each dot represents a single βCDR3aa. The colour scale represents the number of neighbours for each CDR3. Blue dots have only 1 connection while Red dots have more than 3 connections (up to 30).
D. Degree of clustered βCDR3aa. (****p<0.0001, Mann-Whitney test).
E. Generation probability of dispersed and clustered βCDR3aa in DPCD3+ or CD8+ cells. ****p<0.0001, Mann-Whitney test).
F. Correlation between *Pgen* and βCDR3 number of connections in the CD8^+^ thymocyte repertoire. Contour plot represent the generation probability as a function of βCDR3 connections in the CD8^+^ thymocytes for donor P29. Linear regression curves between *Pgen* and number of connections are represented as red dotted lines (“y” represent the regression curve’s equation). Pearson correlation coefficient “R” and p-value “p” are calculated for each individual (Supplementary Figure 3C).
G. PCA analysis of TRB VJ gene combinations in CD8 thymocytes. Blue: dispersed nodes; Red: clustered nodes.

The probability of generation (*Pgen*) of a given TCR varies enormously from one TCR to the other. The clustered TCRs from both DPCD3^+^ and CD8^+^ thymocytes have a significantly higher *Pgen* (p<0.0001) than the dispersed ones (Fig. 1E). *Pgen* also increased significantly (p<0.0001) from DPCD3^+^ to CD8^+^ thymocytes (Fig. 1E). Moreover, there is a robust and remarkable correlation between βCDR3 generation probability and their number of connections (p<0.0001) (Fig. 1F and Fig. S3C).

Clustered βCDR3s belong to TCRs that have a preferential usage of V and J genes, with a significant (p<0.01) overrepresentation of Vβ 12-3, 27, 5-1, and 7-9 and of JB 1-1 and 2-7 (Fig. S3D-E). This results in a clear separation of clustered versus dispersed TCRs in a principal component analysis of VJ usages (Fig. 1G).

### Clustered CDR3s are enriched for publicness

Clustered βCDR3s are also enriched in public sequences, i.e. being shared between at least 2 individuals (p<0.0001) (Fig. 2A). The significant increase of public βCDR3s in CD8^+^ versus DPCD3^+^ thymocytes (p<0.0001) is mostly that of the clustered βCDR3s. For CD8^+^ thymocytes, up to 31.7% of clustered βCDR3s are public compared to barely 1% of the dispersed ones (Fig. 2A, Fig. S4A). Public βCDR3 have a significantly higher generation probability than the private ones (p<0.0001) (Fig. 2B), which is robust across all the patients (Fig. S4B, Table S1). Noteworthily, the number of public βCDR3 shared between individuals is independent of the number of shared HLA Class I alleles (Fig. 2C). Finally, clustered βCDR3s of one individual can be connected to those of other individuals (up to twelve), and more frequently in CD8^+^ versus DPCD3^+^ thymocytes (Fig. 2D and E), indicating a convergence of specificities between individuals’ clustered repertoires. Altogether, these results indicate that the mechanisms for TCR generation and for their further thymic selection are biased to shape a public repertoire of connected βCDR3s with shared recognition properties.

**Figure 2.**
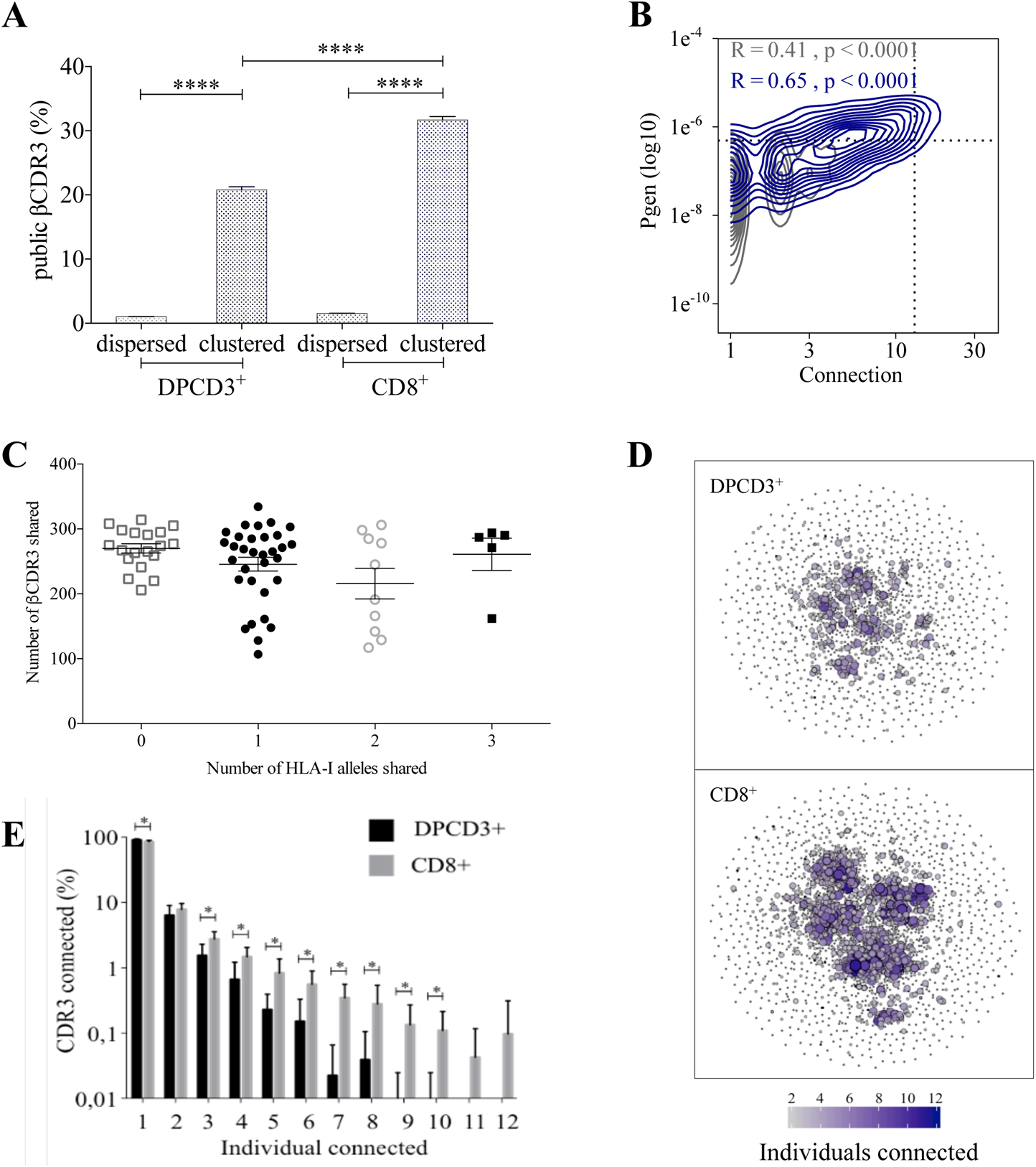
Thymocyte differentiation produces clustered CDR3s with high publicness. A. Mean percentages of public (black) or private (grey) βCDR3s in all, dispersed or clustered nodes. (****p<0.0001, Mann-Whitney test).
B. Enrichment of public βCDR3s in the CD8^+^ thymocyte repertoires. Representation of the generation probability as a function of βCDR3 connections in individuals (Pn). The contour plots represent shared (blue) or private (grey) βCDR3s for a representative patient. The Pearson correlation coefficient “R” and p-value “p” are calculated for each group. The black dotted lines delimit the threshold for the 2.5% sequences with the higher *Pgen* and connection. βCDR3s with both the highest *Pgen* and connections are also the most public for 12 out of 12 individuals (p<0.0001, Chi-square test, Supplementary table S1, Supplementary Figure 4B).
C. The number of public βCDR3s in CD8^+^ thymocytes is independent of the number of HLA-I alleles shared. Each dot represents the number of βCDR3s shared between two donors in the first 18,000 CD8^+^ thymocytes. There is no significant difference in the number of public βCDR3s according to the number of HLA-I alleles shared(p>0.1, Mann-Whitney test). The number of public βCDR3s is independent of the number of HLA alleles shared.
D. Public convergence of βCDR3 recognition properties during thymopoiesis. CDR3 connections between individuals. The top 1,500 βCDR3s were sampled from DPCD3^+^ (upper square) and CD8^+^ (lower square) cells from each individual and pooled. The CDR3s are clustered based on LD≤1 with colour and size both representing the level of sharing between individuals for each CDR3.
E. Convergence of public βCDR3 specificities during thymopoiesis. Bar plots representing the percentage of CDR3s from an individual that are connected to CDR3s of other individuals, for DPCD3^+^ and CD8^+^ thymocytes. The first two bars represent CDR3s that are not connected (n=1). The number of unconnected nodes in DPCD3^+^ is higher than in CD8^+^ (*p=0.002). The other bars represent the percentage of CDR3s connected between individuals. The number of nodes connected to 3 to 10 individuals is significantly higher in CD8^+^ than in DPCD3^+^ cells (*p<0.01, multiple t-test).

### Clustered public CDR3s are enriched in viral specificities

The preferential selection of clustered public CDR3s that could represent over 8% of the sampled repertoire (Fig. 2A) raises the question of their specificities. As the main function of CD8^+^ T cells is cytotoxicity towards virally infected cells, we investigated whether the clustered CDR3s could be associated with virus recognition. We curated two databases of βCDR3s from TCRs specific for human infectious pathogens (*27, 28*) to retain only a set of 5,437 βCDR3 that had been identified by the binding of soluble multimeric MHC/peptide complexes i.e. tetramers or dextramers (*29*). We detected an enrichment of these virus-specific βCDR3s in clustered versus dispersed CD8^+^ thymocytes (p<0.0001) and in CD8^+^ versus DPCD3^+^ thymocytes (p<0.0001) (Fig. 3A). Moreover, these virus-specific βCDR3s were significantly enriched in βCDR3s with the highest *Pgen* and node degree (Fig. 3B and C, Fig. S5A, Table S2). They were also highly shared between individuals (Fig. 3D) and they contained a high representation of CMV, EBV and influenza specificities (Fig. S5B). To better represent the contribution of virus-specific CDR3s to the global repertoire of CD8^+^ cells, we connected them to all CD8^+^ cell CDR3s of the 12 individuals using either a perfect matching (LD=0) (Fig. 3E left panel) or an LD≤1 (Fig. 3E right panel). In the latter representation, the 13,557 connected CDR3s (i) covered most of the clustered CDR3s of all individuals (Fig. 3E, Fig. S6A), (ii) amounted to up to 7.4% and 12.3% of the DPCD3^+^ and CD8^+^ cells CDR3 repertoires, respectively (Fig. 3F), and (iii) were highly shared between individuals (Fig. S6B).

**Figure 3.**
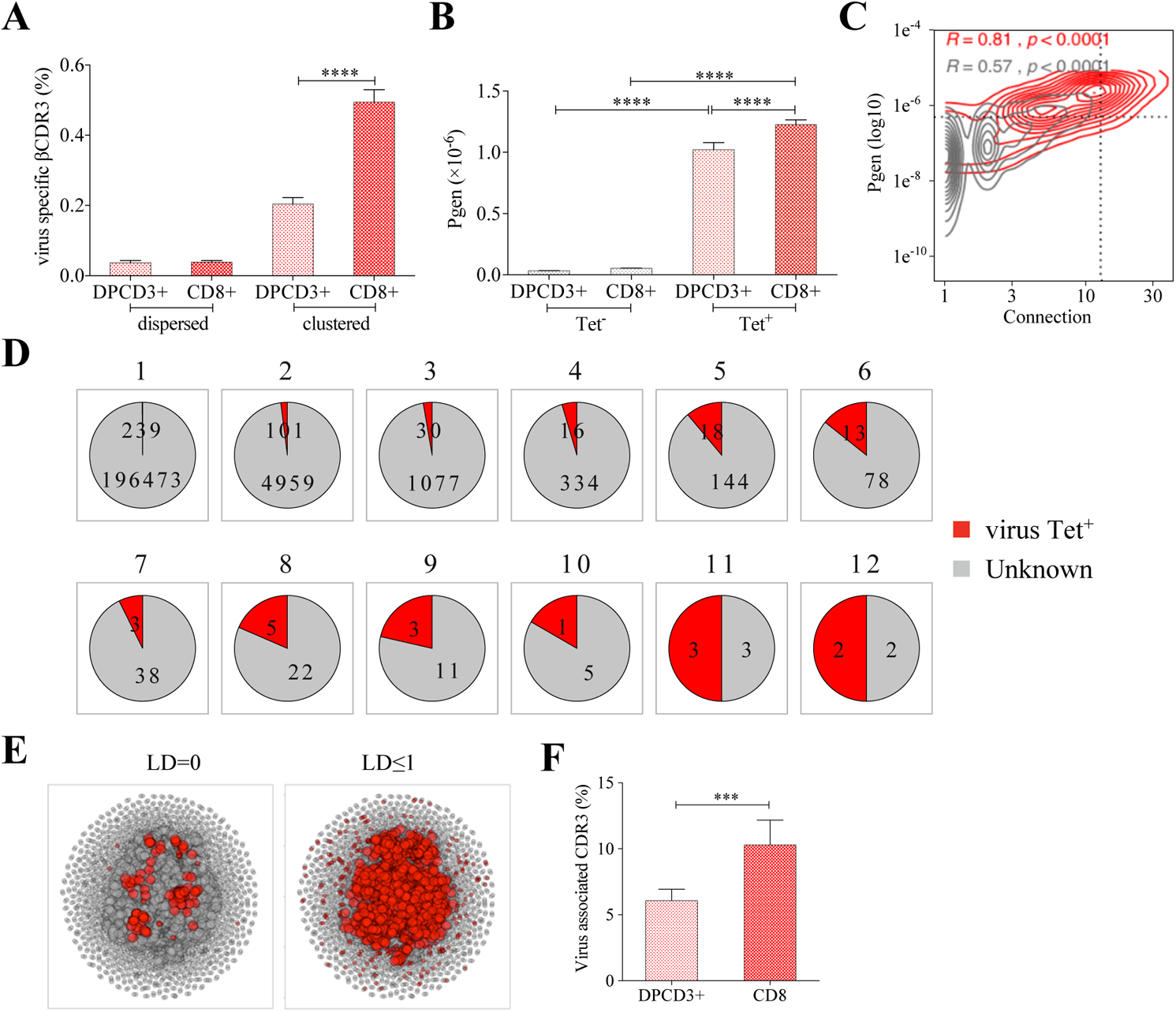
Clustered public TCRs are enriched for virus-specific TCRs. A. Percentages of CDR3s associated with pathogens within DPCD3^+^ and ThyCD8 cells and for dispersed or clustered CDR3s.
B. Mean generation probability of virus-specific βCDR3s, based on their identification by tetramer, in DPCD3^+^ or CD8^+^ thymocytes (****p<0.0001, Mann-Whitney test).
C. Enrichment of virus-specific βCDR3s in the CD8^+^ thymocyte repertoire. Representation of the generation probability as a function of βCDR3 connections in a representative individual. The contour plot represents βCDR3s from TCRs identified as virus-specific based on tetramer identification^12,13^ (red) or with unknown specificity (grey). Pearson correlation coefficient “R” and p-value “p” are calculated for each group. The black dashed lines delimit the threshold for the 2.5% sequences with both higher *Pgen* and degree of connection. (βCDR3s with both the highest *Pgen* and connections were also the most virus-specific for 11 out of 12 individuals, supplementary (p-value <0.0001, Chi-square test).
D. Virus-specific βCDR3s sharing in CD8^+^ thymocytes. Pie charts represent the βCDR3s from private (1) to shared βCDR3s by all donors (12), in grey for βCDR3s with unknown specificity or in red for those with a virus specificity.
E. Network of virus-associated CDR3s. The Shared (identical) and Linked (LD≤ 1) virus-associated CDR3s within the CDR3 network of one individual are in red.
F. Virus antigen coverage. Barplot represents the mean percentage of LD≤ 1 to virus-associated CDR3s in all individuals (***p<0.001, Mann-Whitney test).

Altogether, these results indicate that the selection of clustered CDR3s with high generation probabilities and high inter-individual sharing corresponds, at least in part, to the selection of virus-specific TCRs whose CDR3s are remarkably conserved between individuals independently of their HLA restriction and form what could be referred to as a “paratopic network”.

### Identification of polyspecific TCRs

After thymic selection, an *in-silico* estimation indicated up to 12% of the TCR repertoire as specific for the small set of 30 tested peptides derived from 7 viruses that was used for the analysis (Fig. 3F). This appears hardly compatible with the fact that the TCR repertoire should contain TCRs that react specifically to countless antigens. Actually, we observed that a single cluster of βCDR3s from CD8 thymocytes could comprise unique βCDR3s from TCRs assigned to different viral specificities in independent experiments, for example to CMV, influenza, EBV and YFV (Yellow Fever Virus). This observation is puzzling, as clustered CDR3s are likely to respond to similar antigens (Fig. S1B and (*20–26*)). It suggested that some virus-associated TCRs could have a “fuzzy” specificity that could allow the recognition of peptides from different viruses. We explored the public databases of virus-specific TCRs (*27, 28*) in more detail and discovered that numerous single CDR3s had been assigned to multiple specificities, for example, single CDR3s binding both CMV and influenza tetramers (Fig. S7A). In addition, among the 13,557 virus-associated βCDR3s that we previously estimated in CD8^+^ thymocytes (Fig. 3E, “LD≤1”), almost 7% (958/13557) are *in silico* assigned (LD≤1) as specific for at least two distinct viruses (Fig. S7B).

### Binding properties of polyspecific TCRs

We further investigated the potential T cell specificities of CD8+ thymocytes by using the GLIPH2 algorithm to analyze a combined dataset of 216,000 CDR3β sequences from our CD8^+^ thymocytes and from 32,496 tetramer-specific sequences from the curated VDJdb database. This algorithm groups together CDR3s that have structural characteristics that make them likely to recognize the same antigens, hence called “specificity groups” (*24*). We identified 93,182 such specificity groups. We then analyzed the presence among these groups of TCRs from the VDJdb database with a known viral specificity. Noteworthy, 31.6% of the specificity groups contained at least one CDR3 with a known specificity, an already surprising observation given that the VDJdb database relates to a quite limited sample of the universe of specificities. Even more surprisingly, 6.8% of the clusters identified by GLIPH2 contained CDR3s with more than one specificity, and up to 9 different ones (Fig. 4A).

**Figure 4.**
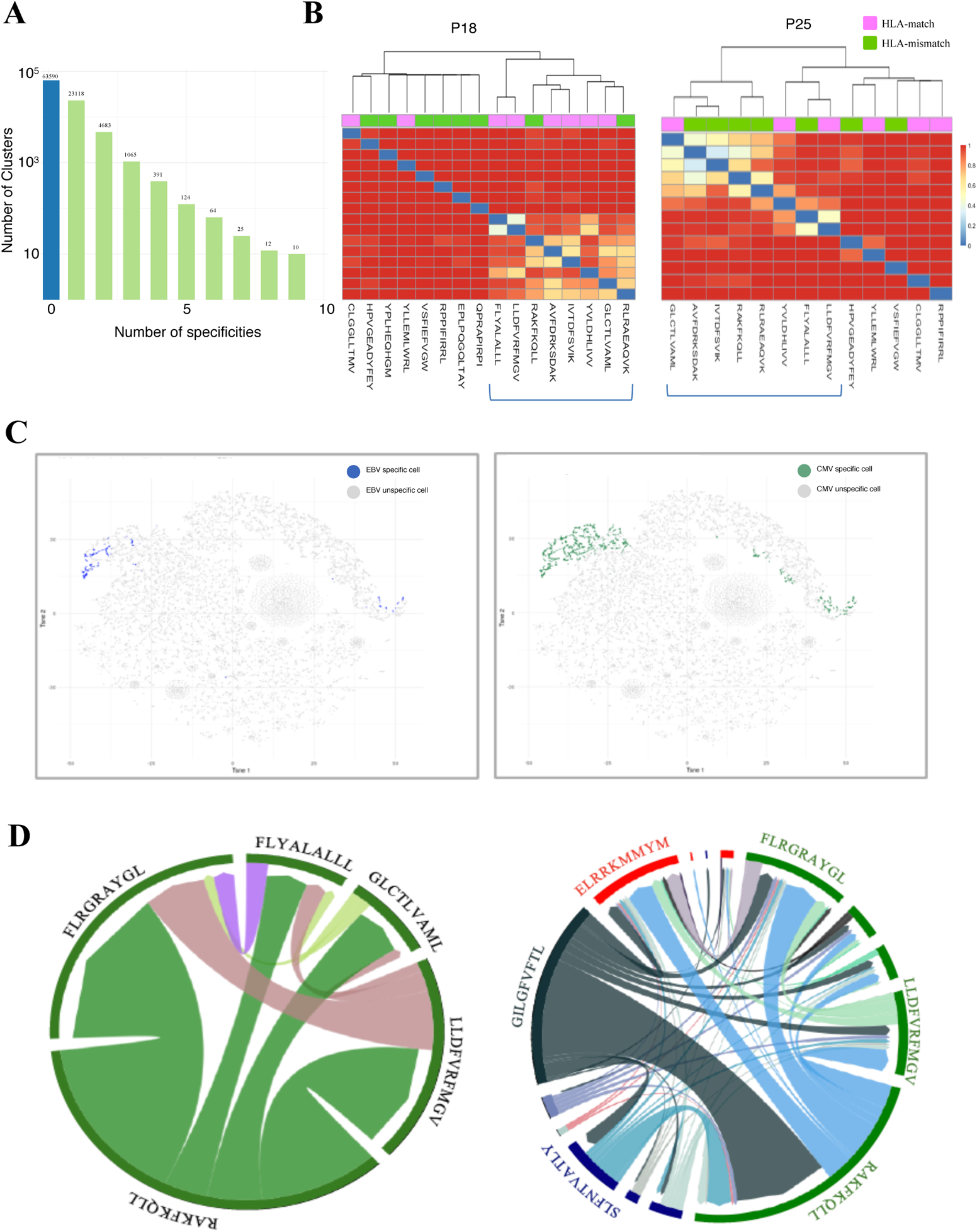
Identification of polyspecific TCRs. A. Analysis of shared T cell specificities with the GLIPH2 algorithm. 216,000 CDR3β sequences from the CD8 thymocytes and 32,496 sequences from VDJdb that are tetramer specific were analysed to obtain 93,182 specificity groups. 31,6% of the groups contained at least one CDR3 with a known specificity for a virus. 6,8% of the clusters identified by GLIPH2 contained CDR3s with more than one specificity, and up to 9 different ones
B. In silico analysis of EBV specific CDR3β. Morisita-Horn dissimilarity analysis of CDR3β repertoire specific for EBV Tetramer identified by GLIPH Algorithm for P18 and P25 patients.
C. t-SNE representation of the single-cell EBV (left panel) and CMV (right panel) TCR specificities from one seronegative individual. CD8^+^ T cells are able to bind both EBV and CMV HLA-matched dextramers. EBV Dex^+^-specific cells are in blue and CMV Dex^+^-specific cells are in green. Some of the cells appear specific for both.
D. Chord diagram showing TCR binding to multiple Dextramers. The chord diagram shows TCR binding to HLA-matched dextramers loaded with peptides from unrelated viruses. Each segment represents TCR binding to the peptides marked above. The size of the segments corresponds to the number of TCRs binding to these peptides. The link between segments identifies multiple TCR binding to different peptides. The colours of the segments represent the different viruses: CMV (red), EBV (green), HIV (dark blue), HPV (light green), HTLV (purple) and influenza (dark grey). The full list of the different peptides is in Supplementary table 3.

We further analyzed the CDR3β repertoires identified with GLIPH2 as specific for at least one of 16 EBV peptides (Fig. 4B). The analysis of these CDR3s based on the Morisita-Horn (MH) dissimilarity index identifies peptides recognized by overlapping repertoires. Noteworthy, the presence of shared CDR3β EBV specific repertoire is observed in both a match/mismatch HLA context. The MDS representation of MH dissimilarity index for EBV repertoire (Fig. S7C) show the convergence of these shared CDR3β EBV specific repertoire across patients.

As these observations are made only from studying unpaired CDR3s, we aimed to confirm them with paired α and β virus-specific CDR3s obtained from single-cell TCR sequencing. These sequences were obtained from 160,914 blood CD8^+^ T cells isolated from four healthy donors and incubated simultaneously with dextramers complexed with peptides from CMV, EBV, HIV, HPV, HTLV and influenza. We could identify numerous single cells harboring TCRs that could bind to multiple dextramers. As an example, single-cell TCRs that bind dextramers matching the HLA of a given individual loaded with EBV or CMV peptides are shown (Fig. 4C). Projecting the binding of EBV dextramers onto a t-SNE representation based on single-cell specificity identified a few distinct regions that contained positive cells (Fig. 4C, left t-SNE). Repeating the same analysis with CMV dextramers identified binding in these same regions, and often on the same cells (Fig. 4C, right t-SNE). Thus, single TCRs appear to bind to both EBV and CMV HLA-matched dextramers.

Actually, unique TCRs could recognize multiple unrelated peptides from the same virus as well as peptides from different viruses. We represented on chord diagram the binding of TCRs to HLA-matched EBV dextramers loaded with different peptides (Fig. 4D left). For example, one half of the TCRs that bind to the RAKFKQLL peptide also bind the FLRGRAYGL peptide. Furthermore, some TCRs recognizing this RAKFKQLL peptide can also bind peptides from CMV (red), EBV (green) or HIV (dark blue) (Fig. 4D right).

Finally, we analyzed in more detail the binding scores of TCRs from 2 patients that all use the same CASSIRSSYEQYF βCDR3 and that were associated with different ⍺CDR3s. This represented 664 cells harboring 55 unique TCRs and 1410 cells harboring 91 unique TCRs, respectively (altogether 2074 cells expressing 131 unique TCRs). We represented the binding score of these CDR3s for 49 different dextramers. Noteworthily, these 2074 TCRs all had the highest binding score for a matched HLA-A*0201 dextramer bound to an influenza peptide (Fig. S8). However, they also had a high binding score for a series of peptides from CMV and EBV bound to HLA-A*0301 and A*1101 dextramers. Most importantly, there was no significant binding to most of the dextramers, indicating that the binding of these few dextramers was specific. These results suggest that polyspecificity is biased towards recognition of common viruses.

### Polyspecific T cells are activated in vitro by multiple viral peptides

These peculiar properties of polyspecific T cell (psT) TCRs led us to assess their functional relevance. We first evaluated the in vitro cross-activation of psT cells with different peptides. Human effector memory CD8^+^ T cells were purified according to their binding of CMV or EBV HLA-matched dextramers (Fig S9); the sorted cells were then stimulated by either the peptide that was used to purify them, or by different ones, and their activation was measured by their IFN-γ production (Fig. 5A). All T cells were efficiently non-specifically activated by PMA/ionomycin. T cells that did not bind any dextramers could not be stimulated by any peptide. In contrast, dextramer-sorted cells could be activated by their cognate peptide and almost as well by the other (Fig. 5B).

**Figure 5.**
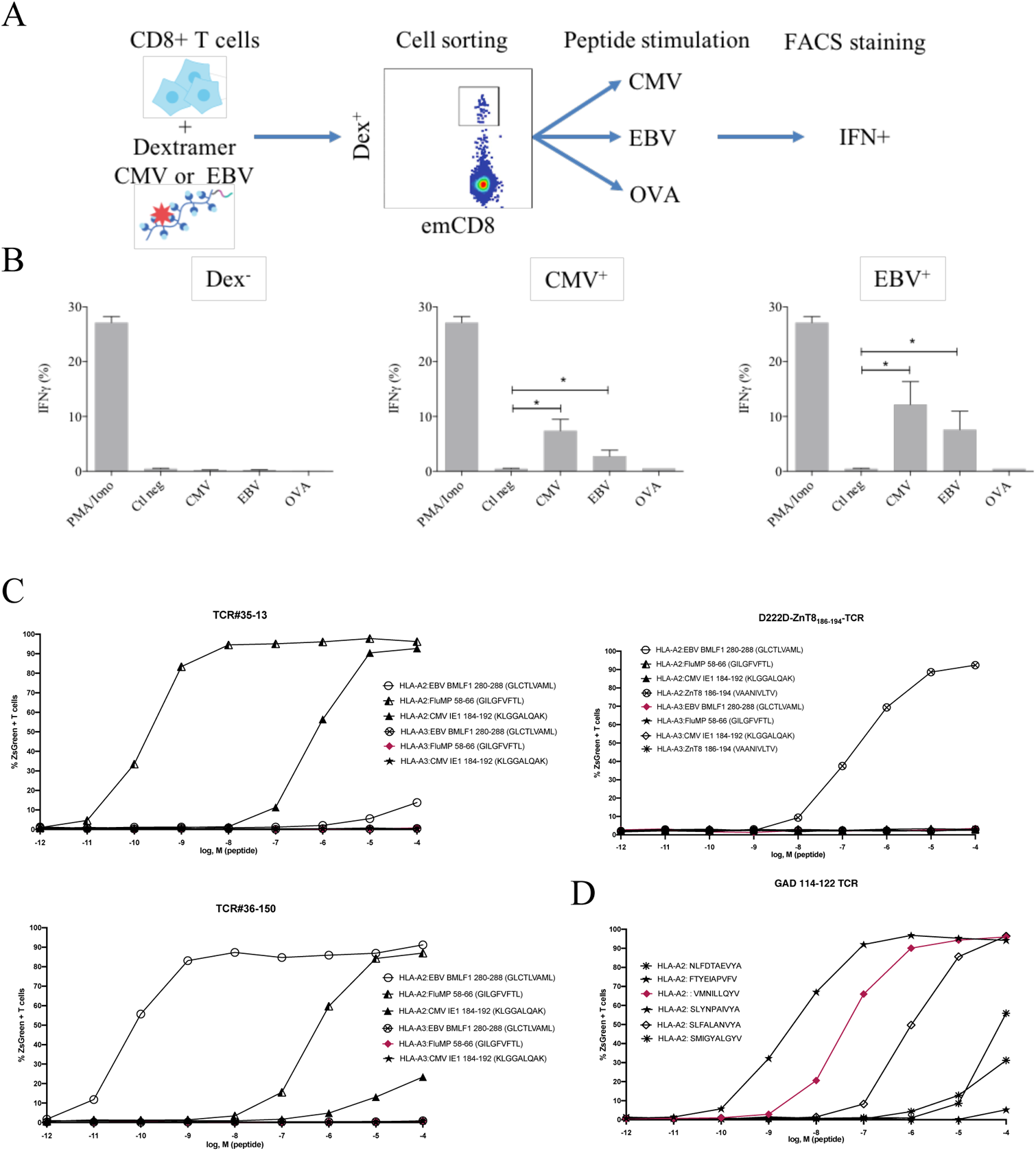
Polyreactivity of polyspecific TCRs. A. Schematic representation of the in vitro cross-activation experiment.
B. In vitro activation of polyspecific T cells. Percentage of IFNγ producing emCD8+ cells after activation with PMA/ionomycin (positive control), no peptide (Ctl neg) or CMV, EBV and OVA peptides (*p<0.05, Mann-Whitney test, mean ± s.e.m.).
C. Poly-reactivity of re-expressed viral epitope-reactive TCRs. 5KC cells transduced with TCR#35-13 (upper left), TCR#36-150 (bottom left) that responded to different peptides in the single cell screening, and D222D-ZnT8186-194 (upper right) that responded to its cognate peptide ZnT8186-194 were stimulated with the indicated nonamer peptides at different concentrations, pulsed on K562 cells expressing either HLA-A*02:01 or HLA-A*03:01. The percent cells expressing the NFAT-driven ZsGreen reporter is shown as activation readout. A representative experiment out of two performed is shown.
D. TCRs from single-sorted CD8+ T cells stained with tetramers loaded with pancreatic self-peptides were cloned in reporter cell lines. We analyzed their response to various peptides: cognate peptide in red and peptides with no significant structural commonalities (grey).

We also tested the response of bulk T cells developed as cell therapy products for the treatment of CMV infections. These cells had been selected through 18 days of stimulation in the presence of a peptide pool covering the complete sequence of the pp65 protein of CMV. Upon stimulation by their cognate peptides, over 85% of the CD8+ T cells produced IFNγ (Fig. S10A). Surprisingly, a small fraction of the same cells produced interferon upon stimulation with unrelated CMV-IE1 peptides, and a large fraction upon stimulation with a peptide pool which covers the sequence of the BZLF-1 protein of EBV (Fig. S10A). There was no interferon production upon stimulation of the same cells by a peptide pool which covers the EBNA1 protein of EBV (Fig. S10A). The stimulation with EBV peptides requires a 100-fold higher but still physiological concentration of peptides (Fig. S10B).

Finally, we directly confirmed the polyspecificity of unique TCRs upon their cloning and re-expression. We used a 5KC murine T-hybridoma cell line devoid of endogenous TCR, and transduced with a ZsGreen fluorescent reporter under the control of nuclear factor of activated T cells (NFAT) and with human CD8 (*30*). These 5KC transductants were then stimulated with K562 cells transduced with either HLA-A*02:01 or HLA-A*03:01 as antigen presenting cells, in the presence of individual peptides at increasing concentrations. From the single cell sequencing dataset, we selected TCRs that could bind dextramers loaded with Flu, CMV and EBV peptides. TCR#35-13 responded strongly to HLA-A*02:01-transduced K562 cells loaded with Flu MP 58-66 (EC50: 2.02×10-9 M) and CMV IE1-184-192 (EC50: 7.99×10-7 M), and poorly to EBV BMLF1 280-288 (EC50: 9.37×10-3 M) (Fig. 5C upper left). TCR#36-150 responded strongly to HLA-A*02:01- transduced K562 cells loaded with EBV BMLF1 280-288 (EC50: 9.99×10-11 M) and Flu MP 58-66 (EC50: 7.57×10-7 M) and poorly to CMV IE1 184-192 (EC50: 2.58×10-3 M) (Fig. 5C bottom left). None of the TCRs responded to peptide-pulsed K562 expressing HLA-A*03:01. In contrast, a control D222D TCR (*31*) known as specific for a ZnT8 186-194 peptide responded to this peptide (EC50: 2.68×10-7 M), but not to the Flu, EBV or CMV peptides (Fig. 5C upper right). We then further analyzed TCRs derived from single-sorted CD8+ T cells stained with tetramers loaded with pancreatic self-peptides. TCRs with known specificities were also cloned and expressed in the reporter cell line. We analyzed their response to a set of peptides comprising their cognate peptide and peptides with no significant structural commonalities, selected by testing combinatorial peptide libraries(*30*). We also observed a marked polyspecificity of these TCRs. As an example, a TCR identified as responding to the VMNILLQYV peptide from the Glutamic Acid Decarboxylase protein responded even better to the structurally unrelated SLYNPAIVYA peptide from human chondroitin sulfate N-acetylgalactosaminyltransferase 2 (Fig 5D).

## DISCUSSION

Our work brings to attention an overlooked phenomenon hidden in the literature. In a simple curation of public datasets of antiviral TCRs that were assigned to respond to known viruses - retaining only those from cells that could bind or respond to a known peptide - we found a surprisingly frequent representation of CDR3s that had been assigned to different viral specificities from independent experiments. Within the 40,939 curated βCDR3, 24% were from TCRs that could bind at least two totally unrelated viral peptides. This observation is highly relevant, as it arises in large part from datasets of full TCRs from single-cell sequencing. Noteworthily, in these latter experiments, cells are co-incubated with multiple dextramers bearing a large set of unrelated peptides; although this dataset is widely available, researchers have missed (or dismissed) that a single cell could bind multiple dextramers (sometimes a dozen). In the methodology description, the technology provider recommends determination of the specificity of a given T cell by the identity of the dextramer that has the highest binding counts on that cell. This is a rather crude readout, as the highest count is often marginally so. Furthermore, it should be realized that, as there are limited numbers of HLA molecules available per cell, there is competition between dextramers for binding at the cell surface. That a single cell binds multiple dextramers bearing unrelated peptides is in our view highly meaningful. As the binding of multimeric HLA/peptide complexes is a paradigmatic method to define specificity, multiple binding indicates multiple specificities. We thus defined TCRs binding multiple HLA/peptide complexes as polyspecific, and by extension called the cells harboring these TCRs as polyspecific T cells (psT cells).

TCRs are notoriously “cross-reactive,” and it is estimated that each TCR could potentially recognize >10^6^ different peptide/MHC combinations (*32–35*). Indeed, the HLA/peptide/TCR *ménage à trois* leaves room for mimotopic cross-reactivity, i.e. the fact that two unrelated peptides can bind differently to a given HLA molecule, but altogether generate a globally similar structure that would be recognized by the same TCR. Also, it was recently shown that peptides need to share only five residues at specific positions to bind the same TCR (*35, 36*). This latter observation has functional relevance as bacterial-derived peptides sharing such 5 residues with a tissue-restricted MOG self-antigen were shown (i) to activate a known MOG-specific TCR as well as a MOG peptide and (ii) to trigger an autoimmune attack against brain tissues upon mouse immunization (*36, 37*). Thus, if many peptides could interact with a given TCR, it is because they share a structural conservation of their TCR interaction surface (*33*).The functional relevance of these observations were described as that “*TCR cross-reactivity enables effective surveillance of diverse self and foreign antigens without necessitating degenerate recognition of non-homologous peptides*”(*33*).

In contrast to this mimotopic cross-reactivity, we did not observe any specific amino acid sharing structure for the multiple peptides recognized by polyspecific TCRs. Thus, polyspecificity appears to be what was described as “*degenerate recognition of non-homologous peptides”* (*33*). We prefer to describe polyspecificity as defining a fuzziness (rather than “degeneration”) in TCR recognition that makes it capable of interacting with multiple unrelated peptides. In this regard, it is noteworthy that B cells have a machinery for somatic mutations of their BCRs that ultimately allows them to generate antibodies with increased affinity (specificity) for antigens. While TCR generation and BCR generation share many common mechanisms, the fact that T cells did not evolve to use such somatic mutations suggest that T cell recognition has been selected to be more fuzzy than stringent.

As TCRs have been essentially selected during thymocyte differentiation for their ability to bind HLA molecules presented by thymic antigen-presenting cells, it could have been that polyspecificity represents some loose remnant of that property. However, it should be emphasized that a single cell does not bind randomly to any homologous-HLA/peptide complexes (Supplementary Fig. 7). Indeed, each polyspecific TCR has its own binding pattern regarding a large set of dextramers, with most that do not bind. Moreover, when two polyspecific TCRs bind to a given dextramer, they have a similar binding score for others that they also bind to. Thus, this binding is not just “*non-specific*” and must obey some structural rules that remain to be defined.

Polyspecific TCR binding properties are distinct from those of innate-like MAIT and NKT cells (*38, 39*). These have a restricted diversity, with an invariant TCRα chain and a constrained TCRβ repertoire, and are MR1- or CD1-restricted, respectively. In contrast, the repertoire of polyspecific TCRs is highly diverse for both the TCR α and β chains, although with a specific usage of VDJ/VJ recombination. Also, polyspecific T cells found in adults cannot be the remnant of fetal/neonatal T cells that have been described as having a promiscuous repertoire, as these cells have repertoires enriched in germline-encoded TCRs(*40, 41*).

The polyspecific nature of these TCRs has a functional relevance, as we show here that cloned polyspecific TCRs indeed trigger cell activation in response to multiple peptides, although with different efficiencies. In agreement with our results, the analysis of the TCR repertoire of lung tumor infiltrating T cells using GLIPH2, as we did here, recently identified a key TCR that, after cloning and expression, could bind and respond to a tumor-specific self-antigen as well as to a peptide from E. coli and one from EBV (*42*). In the same line, a recent study of multiple sclerosis pathogenesis identified autoreactive CD4+ T cell clones that can respond to self-peptides as well as peptides from EBV and from the bacteria *Akkermansis muciniphila*(*43*). This further establishes the existence of psT cells that respond to multiple antigens from different microbes. Also, in the development of the T-scan platform aimed at characterizing the specificity of TCRs(*44*), the authors performed a set-up experiment with CD8 cells activated by the immunodominant NLV peptide from the pp65 protein of CMV. While T-scan could identify two hits related to NLV overlapping peptides, they also detected two hits related to the unrelated CMV I1E CMV protein. Actually, tetramer staining revealed that while 25% of the T cells in the NLV-expanded population bind NLV tetramers, 2% of the T cells bind IE1 tetramers(*44*). Moreover, the fold enrichment of the IE1 specific cells was close to that of NLV specific cells (60 vs 80, respectively). Thus, activation with a single peptide results in a major expansion of cells with TCRs that can also respond to an unrelated peptide.

The fact that CMV, EBV and influenza specificities are highly represented in polyspecific binding patterns likely reflects the biased representation of these specificities in the databases (15.9%, 16.2%, 50.6%, respectively), but also highlights that the detection of the diverse viral specificities of psT cells is largely underestimated due to the very low number of CDR3s with known viral specificities for viruses other than these. Our results led us to estimate that polyspecific TCRs could represent at least 1/5 of the CD8 selected TCR repertoire. Altogether, our results define peculiar binding properties of polyspecific TCRs. This warrants to study their physicochemical characteristics, with a special interest in solving the structure of such TCRs bound to two unrelated peptides.

### Ideas and speculations

Our findings have important implications for the study of the adaptive immune response in health, diseases and immunotherapies. In the field of infectious diseases and vaccination, they prompt reconsideration of the paradigm of highly diverse adaptive immune repertoires providing a highly antigen-specific antiviral immune response. Indeed, there is an intrinsic weakness in the view that a highly diverse TCR repertoire is ideal to protect us from a countless repertoire of constantly evolving viruses. As best illustrated by the deadly outcome of flu in the elderly or young infants, there is a very limited amount of time for the immune system to react against rapidly replicating deadly viruses. For life-threatening situations, the initial recruitment of frequent polyspecific effector T cells might be more efficient and rapid than having to rely on rare cells with stringent specificity that would need a period of expansion to provide enough fighters. In the race between virus replication and the mounting of the immune response, this early response of polyspecific T cells could provide some control of the viral spread that will allow time for the development of an immunologically fittest response to ensure the final control of the infection. This would explain how a very restricted repertoire of only about 1,000 different TCRs arising from a single T cell progenitor was sufficient to cope with viral infections in a child (*11*). They would also explain why children vaccinated against measles in underdeveloped countries have a better life expectancy than unvaccinated ones (when excluding measles-related events), which could be linked to an overall better response to infections with other pathogens (*48, 49*).

While our findings do not challenge that there are specific TCRs nor that there are specific immune responses, they highlight another mechanism of preparedness of the immune system, reminiscent of the role of (i) other unconventional T cells like MAIT and NKT cells(*45*), (ii) TCR activation by bacterial superantigen (*46*) and (iii) natural antibodies specific for microbial determinants (*47*).

Altogether, polyspecific TCRs form a paratopic network, made of TCRs that have a high probability of generation and are positively selected during thymocyte differentiation. These properties indicate that these TCRs have been positively selected during evolution, further supporting their overall beneficial effects. A fuzzy recognition by polyspecific TCRs would explain the so-called “heterologous immunity” (*50, 51*) in which T cell responses to one pathogen can have a major impact on the course and outcome of a subsequent infection with an unrelated pathogen (*17*). We speculate that individual histories of fuzzy immune responses may thus create patient-specific “antigenic sins” that might be responsible for the diverse quality of immune responses to viruses, from inapparent infection to fulminant immunopathology (*52–54*). Further studies will have to evaluate the contribution of fuzzy immune responses to the efficacy but also the immunopathology of antimicrobial responses and to autoimmunity. We speculate that the origin of many immune diseases should be sought in the repeated activation of polyspecific T cells.

## MATERIALS AND METHODS

### Patients and samples

Thirteen human thymus samples were obtained from organ donors undergoing surgery (Department of Cardiac Surgery, Pitié-Salpêtrière Hospital, France) after approval by the *Agence de Biomédecine* and the *Ministry of Research*. Their age at the time of sampling ranged from 19 to 65 years old. The male-to-female sex ratio was 2.6.

For cross-activation experiments, six leukapheresis samples were freshly collected from healthy donors at EFS Paris Saint-Antoine-Crozatier (Etablissement Français du Sang, Paris, France) after informed consent and according to institutional guidelines. Donor selection was based on matching HLA-A2 class I allele.

### Isolation of thymocytes and extraction of RNA

Single-cell suspensions were prepared from the thymus by mechanical disruption through nylon mesh (cell strainer). Single-cell suspensions from whole thymus were stained with antibodies anti-CD3 (AF700), anti-CD4 (APC), anti-CD8 (FITC). Cells were sorted by fluorescent activated cell sorting (Becton Dickinson^TM^ FACSAria II) with purity >95% to collect populations based on the following labeling: DPCD3^−^ were gated as CD3^−^CD4^+^CD8^+^, DPCD3^+^ were gated as CD3^+^CD4^+^CD8^+^ and CD8^+^ were gated as CD3^+^CD4^−^CD8^+^. RNA was isolated from sorted populations by means of lysis buffer with the RNAqueous-Kit (Invitrogen®) extraction kit, according to the manufacturer’s protocol.

### TCR repertoire library preparation and sequencing

TCR repertoire library preparation and sequencing was performed has previously described(*55*). Briefly, T cell receptor (TCR) alfa and beta libraries were prepared on 100 ng of RNA from each sample with the SMARTer Human TCR a/b Profiling Kit (Takarabio®) following the provider’s protocol. Briefly, the reverse transcription was performed using TRBC reverse primers and further extended with a template-switching oligonucleotide (SMART-Seq® v4). cDNAs were then amplified following two semi-nested PCRs: a first PCR with TRBC and TRAC reverse primers as well as a forward primer hybridizing to the SMART-Seqv4 sequence added by template-switching and a second PCR targeting the PCR1 amplicons with reverse and forward primers including Illumina Indexes allowing for sample barcoding. PCR2 were then purified using AMPure beads (Beckman-Coulter®). The cDNA samples were quantified and their integrity was checked using DNA electrophoresis performed on an Agilent 2100 Bioanalyzer System in combination with the Agilent DNA 1000 kit, according to the manufacturer’s protocol. Sequencing was performed with Hiseq 2500 (Illumina®) SR-300 protocols using the LIGAN-PM Genomics platform (Lille, France).

### TCR deep sequencing data processing

FASTQ raw data files were processed for TRA and TRB sequences annotation using MiXCR (*56*) software (v2.1.10) with RNA-Seq parameters. MiXCR extracts TRAs and TRBs and provides corrections of PCR and sequencing errors.

### Network generation and representation

To construct a network, we computed a distance matrix of pairwise Levenshtein distances between CDR3s using the “stringdist”(*57*) R package. When two sequences were similar under the defined threshold, LD>1 (i.e., at most one amino acid difference), they were connected and designated as “clustered” nodes. CDR3s with more than one amino acid difference from any other sequences are not connected and were designated as “dispersed” nodes.

Layout of networks for fig. 1C, 2D were obtained by using the graphopt algorithm of the “Igraph”(*58*) R package and plotted in 2D with “ggplot2” to generate figures (*59*). Only clustered nodes are represented, edges are not shown and colors represent the node degree (log scale). Layouts of detailed networks in fig. 1A were done with Cytoscape(*60*).

### Statistical analysis and visualization

Normalization was performed by sampling on the top α or β 18,000 CDR3s based on their frequency in each sample. The repertoires with less than 18,000 α or β CDR3s were not included in the statistical analysis. The numbers of samples included in the statistical analysis for the β repertoire were: two for DPCD3^−^, ten for DPCD3^+^ and twelve for CD8^+^. The numbers of samples included in the statistical analysis of the α repertoire were: six for DPCD3^+^ and ten for CD8^+^. Statistical tests used to analyze data are included in the figure legends. Comparisons of two groups were done using the Mann-Whitney test (fig.1B-D-E, fig.2A-C, fig.3A-B-F) and multiple t-test (fig. 2E). The correlation coefficient was calculated using the Pearson correlation coefficient (fig. 2B & 2C, fig. 2C). Enrichment of public CDR3s or virus-associated CDR3s was done using the two-tailed Chi-square test with Yates correction (Tables S1, S2 & S3). Statistical comparisons and multivariate analyses were performed using Prism (GraphPad Software, La Jolla, CA) and using R software version 3.5.0 (www.r-project.org). PCA was performed on the frequency of VJ combination usage frequency within each donor using the factoextra R package. T-SNE were generated using the binding scores of each cell across all the antigens present in the dataset. The function Rtsne of the homonymous R package (*61*) was applied with the perplexity parameter set to 10.

### Probability of generation calculation

The generation probability (*Pgen*) of a sequence is inferred using the Olga algorithm, which is inferred by IGoR (*63*), for fig. 1E and fig.2B. IGoR uses out-of-frame sequence information to infer patient-dependent models of VDJ recombination, effectively bypassing selection. From these models, the probability of a given recombination scenario can be computed. The generation probability of a sequence is then obtained by summing over all the scenarios that are compatible with it. We also used OLGA to generate a random repertoire of 500,000 sequences for each α or β repertoire and 3 down sampling (of unique sequences) to get a control repertoire equal to the size of COVID-19 BAL dataset used in fig. 7B & C. The control repertoire was parametrized by the predefined genomic templates provided with the package.

### CDR3 connections between individuals

In fig. 2D, the top 1,500 βCDR3s were sampled from each of the 12 datasets of DPCD3^+^ and CD8^+^, then merged to obtain two datasets of 18,000 βCDR3s for DPCD3^+^ and CD8^+^. We generated and represented networks, as described above, to investigate the βCDR3 inter-individual network structure.

### Virus-specific CDR3 tetramer public databases

The virus-associated CDR3 databases used for the search for specificity was compiled from the most complete previously published McPAS-TCR (*27*) and VDJdb (*28*) databases. Virus-associated βCDR3s were selected from the original datasets only when derived from a TCR of sorted CD8 T cells that were bound by a specific tetramer. A total of 5,437 such unique tetramer-associated βCDR3s were identified and used. Peptides used for tetramer sorting were from cytomegalovirus (CMV), Epstein-Barr virus (EBV), hepatitis C virus (HCV), herpes simplex virus 2 (HSV2), human immunodeficiency virus (HIV), influenza and yellow fever virus (YFV).

### Virus-specific CDR3 single-cell dextramer public dataset

This dataset contains single-cell alpha/beta TCRs from 160,914 CD8^+^ T cells isolated from peripheral blood mononuclear cells (PBMCs) from 4 healthy donors. Briefly, 30 dCODE™ Dextramer® reagents (Immudex®) with antigenic peptides derived from infectious diseases (9 from CMV, 12 from EBV, 1 for influenza, 1 for HTLV, 2 for HPV and 5 for HIV) were simultaneously used to mark cells. Each Dextramer® reagent included a distinct nucleic acid barcode. A panel of fluorescently labeled antibodies was used to sort pure Dextramer®-positive cells within the CD8^+^ T cell population using an MA900 Multi-Application Cell Sorter (Sony Biotechnology) in a reaction mix containing RT Reagent Mix and Poly dT RT primers. The Chromium Single Cell V(D)J workflow generates single cell V(D)J and Dextramer® libraries from amplified DNA derived from Dextramer®-conjugated barcode oligonucleotides, which are bound to TCRs. Chromium Single Cell V(D)J enriched libraries and cell surface protein libraries were quantified, normalized, and sequenced according to the user guide for Chromium Single Cell V(D)J reagent kits with feature barcoding technology for cell surface protein. We used this dataset to study the presence of multiple specificities in TCR and CDR3. There were 139,378 unambiguous TCRs (with only one α and one β chain). We set the threshold defining positive binding at UMI counts greater than 10 for any given dextramer. This identified 15,195 unique virus-specific TCRs with at least one binding.

### Cross-activation experiment

PBMCs were separated on Ficoll gradient. CD8+ T cells were isolated from PBMCs by positive isolation using the DYNABEADS® CD8 Positive Isolation Kit (Thermo Fisher Scientific) according to the manufacturer’s instructions. emCD8 T cells were purified after staining with CD3-AF700, CD8-KO, CD45RA-PeCy7 according to the manufacturer’s instructions. The samples were also stained either with CMV pp65 NLVPMVATV or with EBV BMLF-1 GLCTLVAML PE-conjugated Dextramers (Immudex®). emCD8^+^Dex^+^ cells were sorted by FACS (FACS Aria II®; BD Biosciences) with a purity >95%. Sorted cells were cultured at a maximum of 5 × 10^5^ cells/mL in round-bottom 96-well plates in RPMI 1640 medium supplemented with 10% FCS, 1% penicillin/streptomycin and glutamate at 37°C with 5% CO_2_. *In vitro* stimulation was performed 24 hours after cell sorting. Sorted cells were stimulated for 6 hours with either nothing or 1 μg/mL of SIINFEKL ovalbumin peptide (OVA), NLVPMVATV cytomegalovirus pp65 peptide (CMV) or GLCTLVAML Epstein-Barr virus BMLF-1 peptide (Ozyme®). The positive control (Ctl PMA/Iono) was performed with 50 ng/mL phorbol myristate acetate (PMA) and 1 mM ionomycin. Intracellular IFN-γ production with an IFN-γ-FITC antibody (BD Pharmingen) was detected in the presence of Golgi-Plug (BD Pharmingen®) after fixation and permeabilization (BD Cytofix/Cytoperm). Data were acquired using a Navios flow cytometer and analyzed with Kaluza analysis software (Beckman Coulter).

### In vitro CMVpp65 CTL selection

Peripheral blood mononuclear cells (PBMCs) were obtained from blood donors at the Etablissement Français du Sang (EFS) with informed consent (Blood products transfer agreement relating to biomedical research protocol 97/5-B—DAF 03/4868). CMV+ donor PBMCs were resuspended in the serum-free culture medium TexMacs (Miltenyi-Biotech) at 10^7^ cells/mL and at a density of 5 × 10^6^ cells/cm^2^. Then, 20 µl PepTivator-CMVpp65 (Miltenyi-Biotech (130-093-435), 30 nmol of each peptide and approximately 140 peptides were added per mL of cell suspension. The final concentration of PepTivator-CMVpp65 was 0.6 nmol (approximately 1 µg of each peptide/mL). With these stimulation conditions, PBMCs were incubated for 4 hours and then transferred to culture flasks at 10^6^ cells/mL in RPMI medium with 8% human serum (EFS Pays de la Loire, France) and 50 IU IL-2/mL. CTL were maintained in culture for 25 days.

### Detection of CMV pp65-specific T cells by intracellular staining with anti-INF-γ-PE

CMVpp65 selected CTL were stimulated for 6 hours in TexMacs medium (without human serum and IL-2) with 20 µL PepTivator-CMVpp65 per mL of cell suspension. Brefeldin-A was added at 10 µg/ml. Negative control (CTL without stimulation) and positive control (CTL stimulated with PMA-ionomycin) were also included. Stimulation with irrelevant peptides was also performed with PepTivator-CMV IE-1 (Miltenyi (130-093-494) and PepTivator-EBV BZLF1 (Miltenyi (130-093-612). Cells were then stained with anti-CD8-FITC, fixed, permeabilized and intracellularly stained with anti-INF-γ-PE.

### Peptide dose-response of TCR transductants

TCR transductants were generated as described [Mann, Zhou, Front Immunol 2020]. Briefly, 5KC T-hybridoma cells (kindly provided by M. Nakayama, Barbara Davis Center, Aurora, CO) were transduced with the NFAT-driven fluorescent reporter ZsGreen-1 along with human CD8 by spinoculation with retroviral supernatant produced from phoenix-eco cells (ATCC CRL-3214). These cells were subsequently transduced with retroviral vectors encoding a chimeric TCR alpha gene followed by a porcine teschovirus-1 2A (P2A) peptide and a chimeric TCR beta gene synthesized by TwistBioscience. K562 cells (ATCC CCL-243) were transduced with lentiviral vectors (TakaraBio) encoding HLA-A*02:01 or HLA-A*03:01 by spinoculation with ultracentrifuge concentrated viral particles produced from 293T-HEK-TN cells (System biosciences), followed by sorting of cells stained with anti-human HLA-A, B, C antibody conjugated with PE-Cy7 (clone W6/32, BioLegend).

TCR transductants (20,000 cells per well) were stimulated for 18 h with individual peptides at different concentrations (9 10-fold serial dilutions from 100 μM) in the presence of HLA-A*02:01+ or HLA-A*03:01+ K562 cells (50,000 cells per well), followed by analysis of ZsGreen-1 expression. Peptides (from Synpeptide) tested for the response by TCRs are GILGFVFTL (Flu MP 58–66), GLCTLVAML (EBV BMLF1 280-288), KLGGALQAK (CMV IE1 280-288), VMNILLQYV (GAD65 114-122), VAANIVLTV (ZnT8 186-194) and other sequences selected by testing a GAD 114-122-reactive TCR by combinatorial peptide libraries, as described(*30*) EC50 values were calculated using the nonlinear regression log (agonist) vs. response (three parameters) equation model in GraphPad Prism 9.

## Supplementary Materials

**Fig. S1.** Thymocytes cell sorting dot plot.

**Fig. S2.** ßCDR3 network during thymopoiesis.

**Fig. S3.** Properties of thymocytes’ TCRs.

**Fig. S4.** Publicness of thymocytes’ CDR3s.

**Fig. S5.** Virus specificities of thymocytes’ CDR3s.

**Fig. S6.** Sharing of virus specific thymocytes’ CDR3s.

**Fig. S7.** Binding properties of single cell TCRs.

**Fig. S8.** βCDR3s drive polyspecificity.

**Fig. S9.** Gating strategy for sorting effector memory (emCD8) Dextramer positive cells.

**Fig. S10.** Polyspecific properties of anti-CMV cytotoxic cells.

**Table S1.** Enrichment of public **β**CDR3s in CD8+ thymocytes vs DPCD3+.

**Table S2.** Enrichment of virus-specific **β**CDR3s from databases^14,15^ in clustered CD8+ thymocytes.

**Table S3.** List of peptides represented on the chord plot. Related to 4I.

## Acknowledgments

The authors would like to express their gratitude to the organ donors and their families who allowed the collection of samples for research under sad circumstances. The authors would also like to thank Prof. Pascal Leprince, Dr. Guillaume Lebreton, and Dr. Marina Rigolet of the cardiac surgery team, Prof. Bruno Riou and the graft coordination team of the Pitié-Salpêtrière hospital for their invaluable contribution to sample collection. The authors thank the UMR 8199 LIGAN-PM Genomics platform (Lille, France) for efficient sequencing. We also thank Thierry Mora, and Aleksandra Walczak for helpful discussion.

## Funding

This work was primarily funded by:

TRiPoD ERC-Advanced EU grant 322856 (DK).

LabEx Transimmunom ANR-11-IDEX-0004-02 (DK).

RHU iMAP ANR-16-RHUS-0001 (DK).

Besides, ZZ was supported by the JDRF Postdoctoral Fellowship 3-PDF-2020-942-A-N and RM by the Agence Nationale de la Recherche ANR-19-CE15-0014-01 and Fondation pour la Recherche Médicale EQU20193007831.

## Author contributions

Conceptualization: DK.

Data curation: VQ, PB, PS, HPP.

Data Analyses: VQ, PB, FM, VM, PS, HV, ZZ, HPP.

Funding acquisition: DK

Investigation: VQ, PB, VM, PS, HV, FM, NC, MBS, HPP, BC, HV, ZZ, BB, MS, EMF.

Methodology: VQ, DK.

Project administration: VQ, DK. Resources: VQ, PB.

Software: VQ, PB, PS, HPP.

Supervision: DK.

Validation: DK.

Visualization: VQ, DK.

Writing – original draft: VQ, DK.

Writing – review & editing: VQ, DK.

## Data and materials availability

Datasets from VDJdb were downloaded from https://vdjdb.cdr3.net. Datasets from McPAS-TCR were downloaded from http://friedmanlab.weizmann.ac.il/McPAS-TCR/. We manually curated these datasets to be sure to use only βCDR3s from CD8 tetramer-specific cells.

Single-cell datasets from 10X genomics were downloaded from https://support.10xgenomics.com/single-cell-vdj/datasets (‘Application Note - A New Way of Exploring Immunity’ section, datasets ‘CD8+ T cells of Healthy Donor’ 1–4, available under the Creative Commons Attribution license).

Data from the donors are available on request to the authors.

---

**Figure S1:**
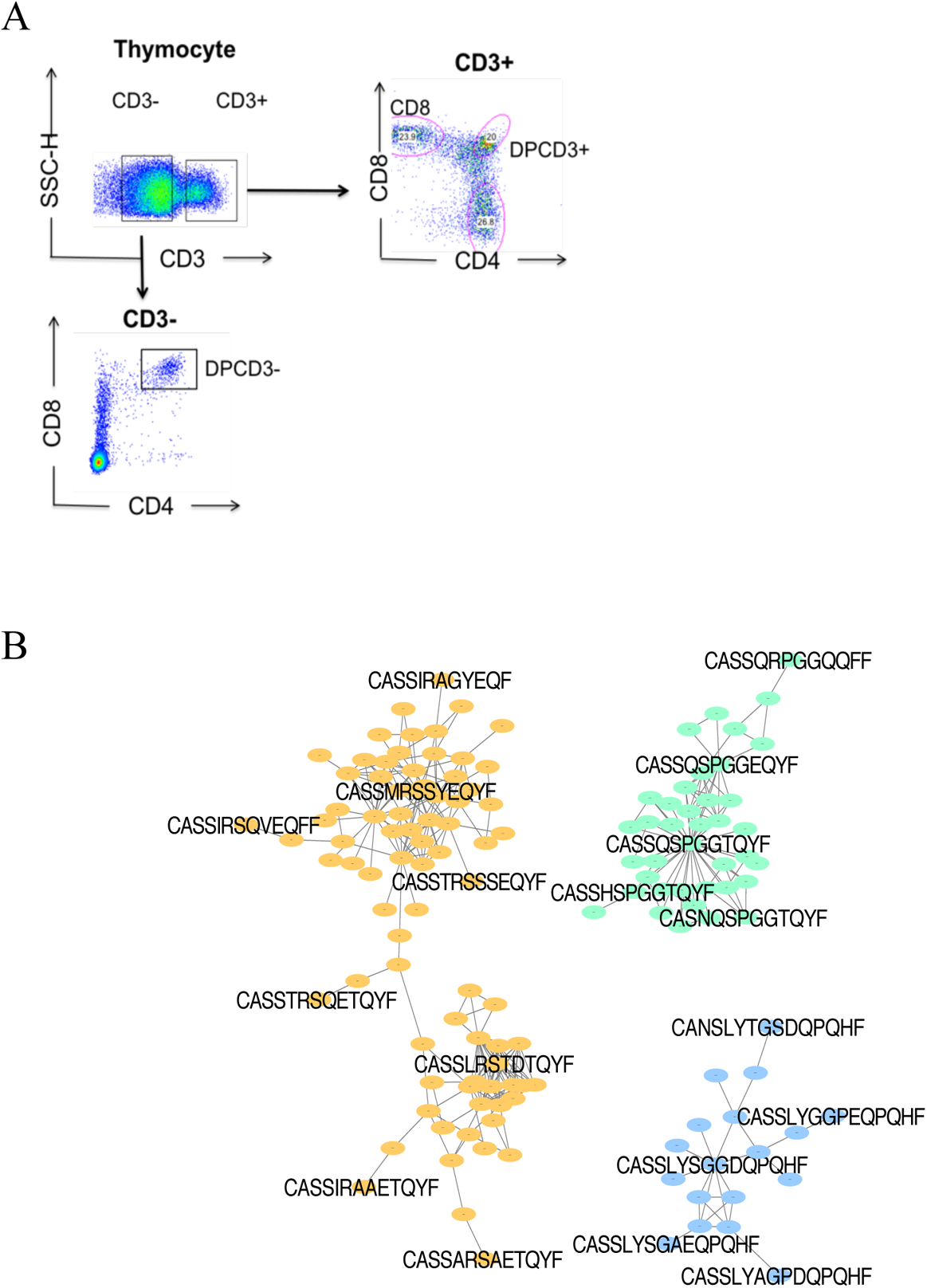
Dotplot of thymocytes sorting. A. Dotplot of thymocytes sorting. Cells from the different patients were sorted according to their phenotypes: CD3+CD4-CD8+ for ThyCD8, CD3+CD4+CD8+ for DPCD3+ and CD3-CD4+CD8+ for DPCD3-.
B. Networks of βCDR3s specific for GILGFVFTL from influenza (orange), GLCTLVAML from Epstein-Barr virus (green) and FPRPWLHGL from human immunodeficiency virus (blue) are shown. These βCDR3s are from TCRs identified on CD8 T lymphocytes isolated with class I tetramer loaded with the indicated peptides(*27, 28*). Each node represents a clonotype. Two different clonotypes are connected if their βCDR3s differ by at most one amino acid (LD≤1).

**Figure S2:**
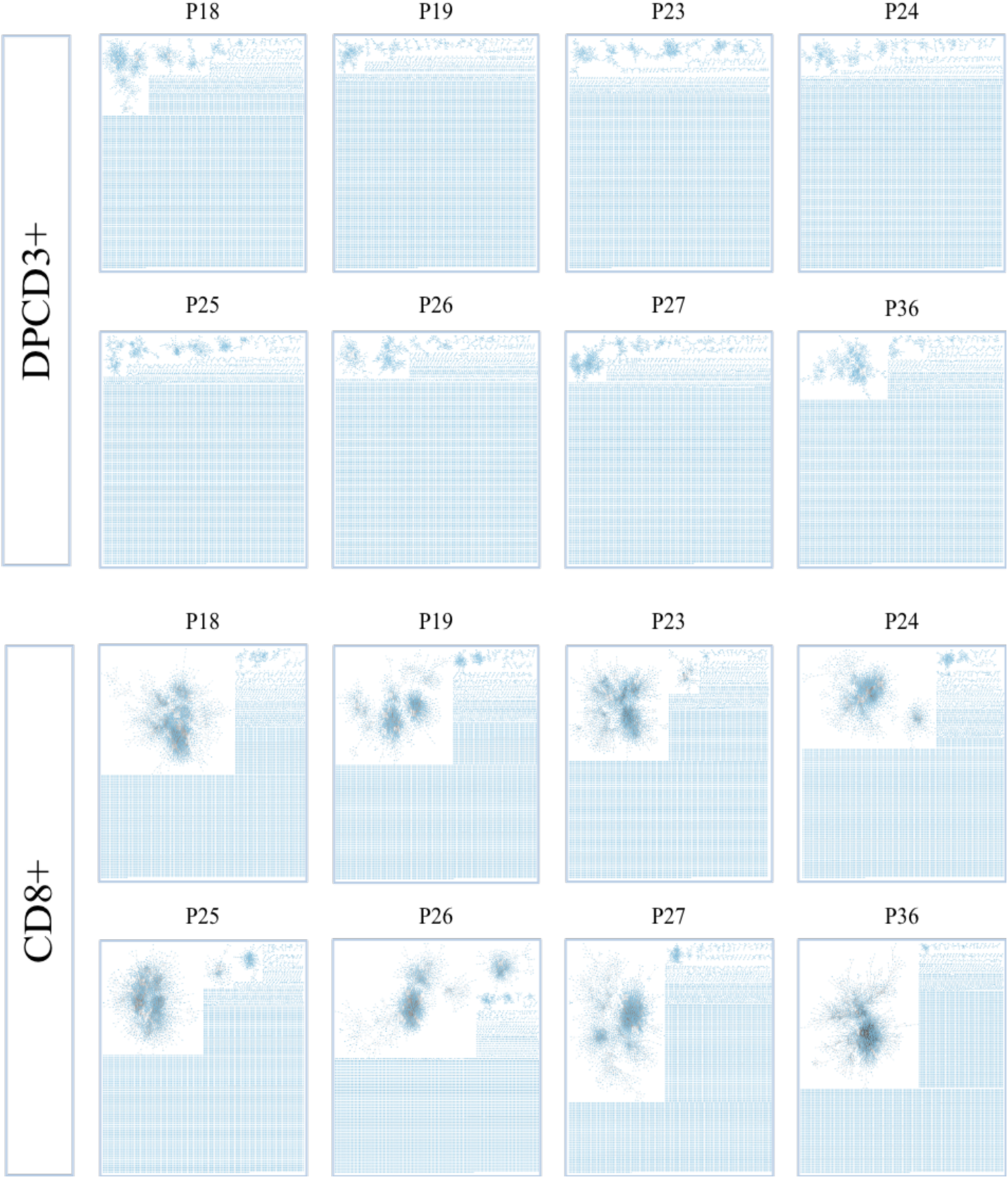
Properties of thymocytes’ TCRs. βCDR3 network during thymopoiesis. Representation of the 18,000 most frequent βCDR3 networks from DPCD3^+^ and CD8^+^ thymocytes of eight donors (Pn).

**Supplementary Figure 3.**
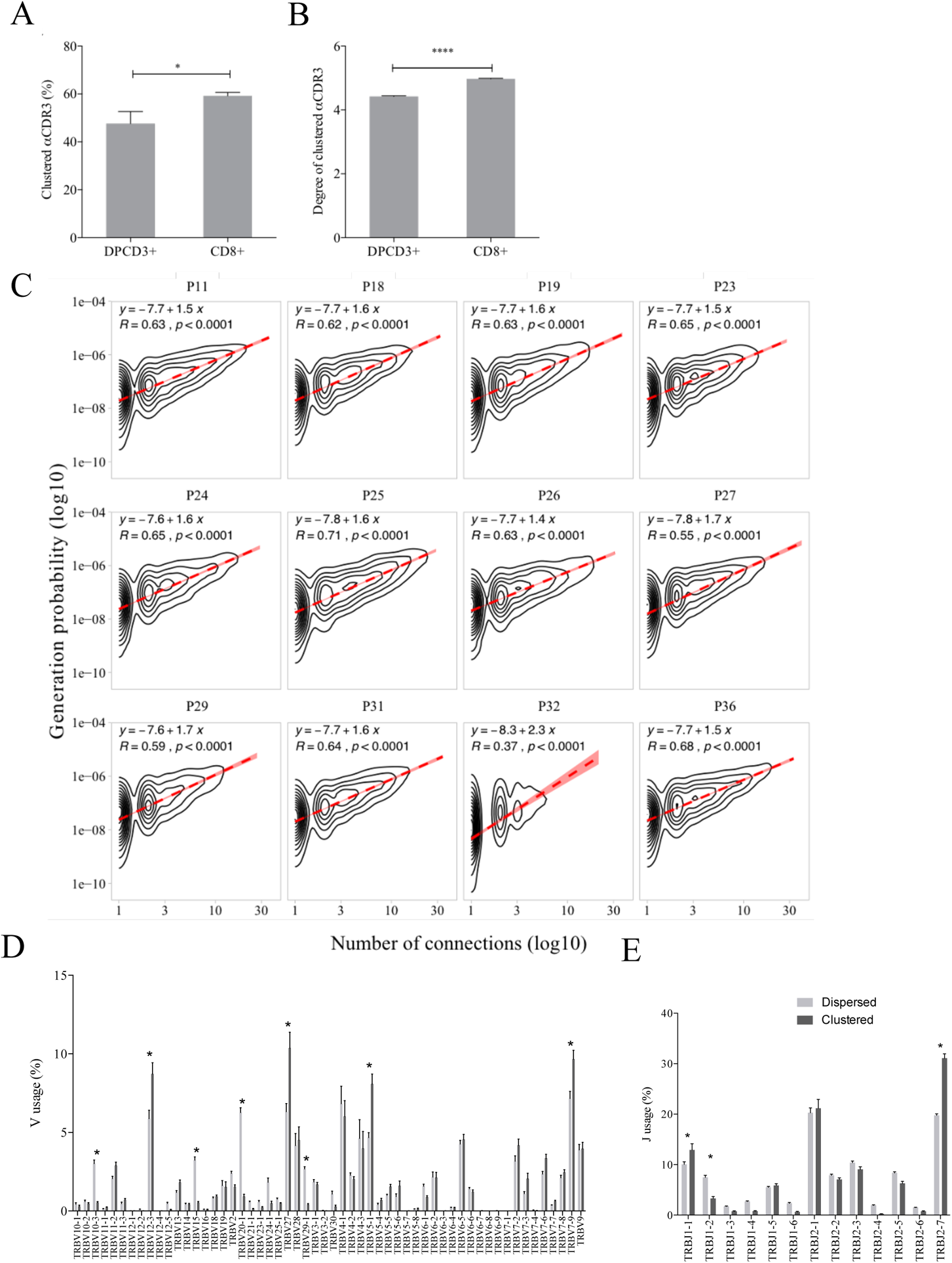
(A-B) Clustered αCDR3s from DPCD3^+^ and CD8^+^ thymocytes. Analyses were performed on the first 18,000 most frequent αCDR3s per sample (n=6 for DPCD3^+^ and n=10 for CD8^+^ thymocytes). (A) Percentage of clustered αCDR3s. (mean±s.e.m., *p=0.016, Mann-Whitney test). (B) Degree of clustered αCDR3s. (mean±s.e.m., ****p<0.0001, Mann-Whitney test). (C) Correlation between *Pgen* and βCDR3 number of connections in the CD8^+^ thymocyte repertoire. The contour plots represent the generation probability as a function of βCDR3 connections in the CD8^+^ thymocytes for donors P11 to P36. Linear regression curves between *Pgen* and number of connections are represented as red dashed lines (“y” represents the regression curve’s equation). The Pearson correlation coefficient “R” and p-value “p” are calculated for each individual. (D-E) Clonogram representation of TCR Vβ (E) and Jβ (F) usage in clustered versus dispersed CD8^+^ thymocytes. The bar plots represent the mean percentage of TCR Vβ (left) and Jβ (right) in dispersed (light grey) versus clustered (dark grey). (* p<0.01, multiple t-test).

**Figure S4.**
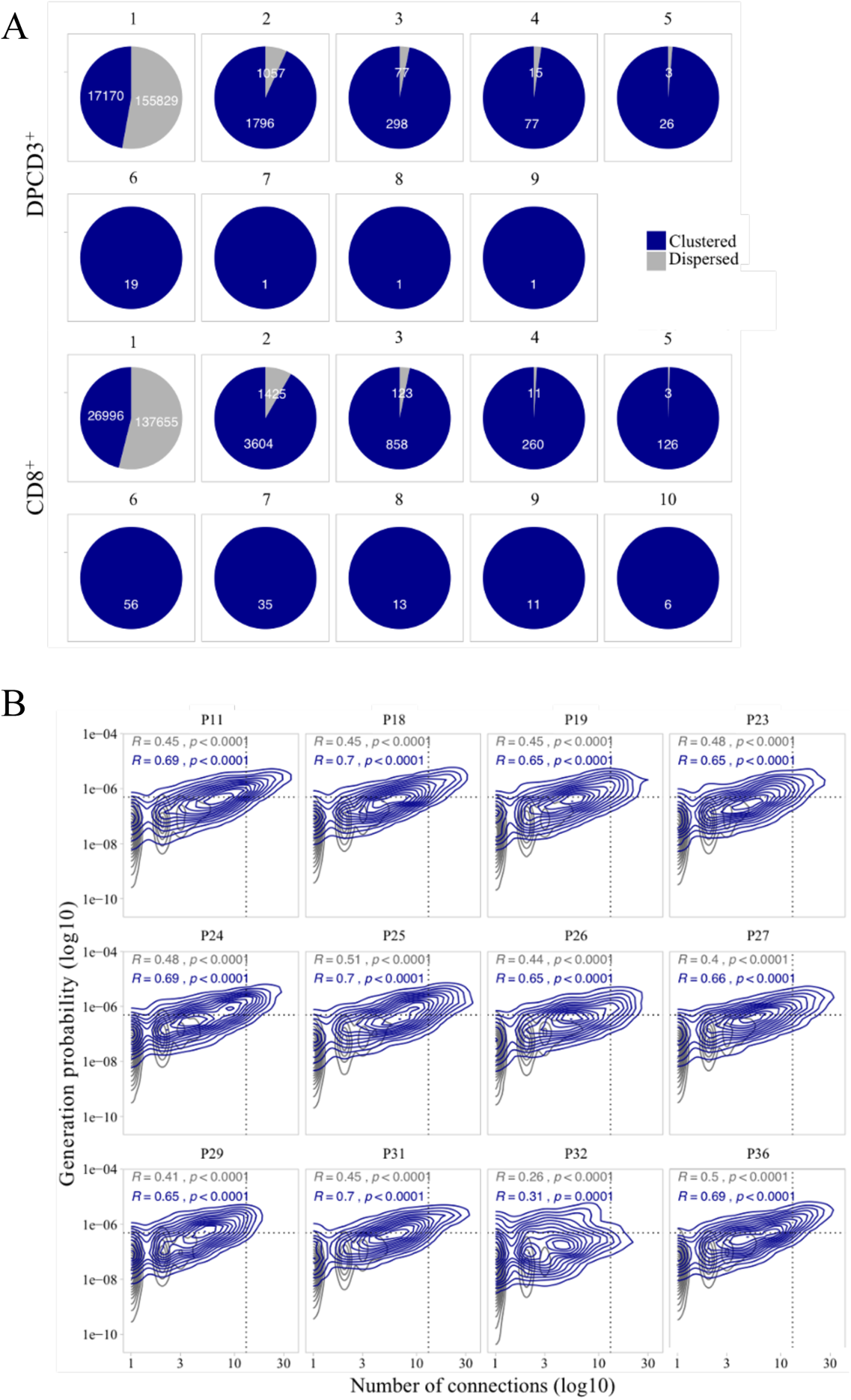
Publicness of thymocytes’ CDR3s. (A) βCDR3 sharing between individuals. Pie charts represent the sharing between individuals before (DPCD3^+^) and after thymic selection (CD8^+^). Colours represent the dispersed (grey) or clustered (blue) CDR3s. Sharing was analyzed within the 10 donors for which there were at least 18,000 βCDR3s in DPCD3^+^ and in CD8^+^ thymocytes. (B) Enrichment of public lllCDR3s in the CD8+ thymocyte repertoires. Representation of the generation probability as a function of lllCDR3 connections in individuals (Pn). The contour plots represent shared (blue) or private (grey) lllCDR3s. The Pearson correlation coefficient “R” and p-value “p” are calculated for each group. The black dotted lines delimit the threshold for the 2.5% sequences with the higher Pgen and connection. lllCDR3s with both the highest Pgen and connections are also the most public for 12 out of 12 individuals (p<0.0001, two-tailed Fisher test).

**Figure S5.**
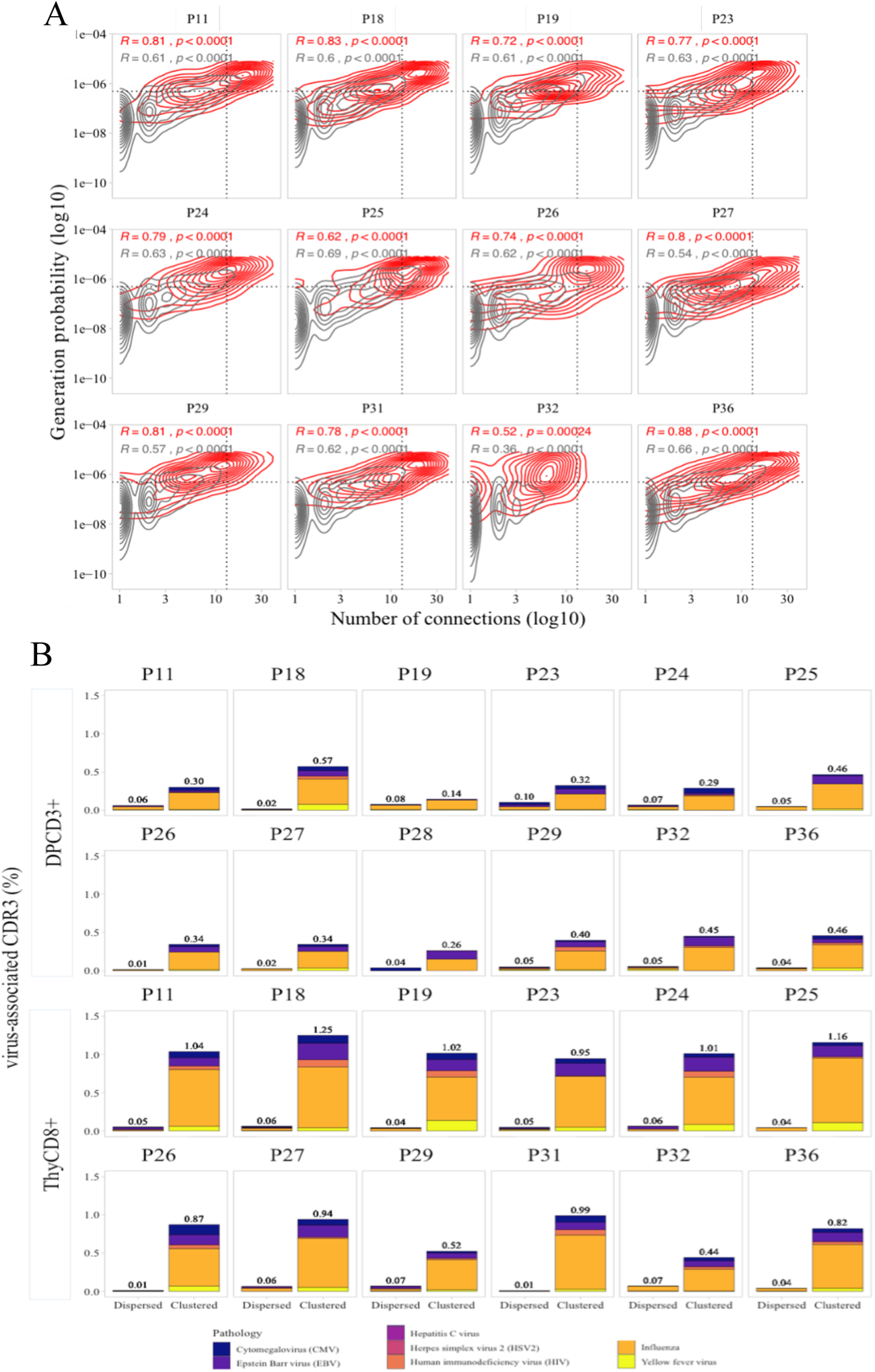
Virus specificities of thymocytes’ CDR3s. (A) Enrichment of virus-specific lllCDR3s in the CD8+ thymocyte repertoire. Representation of the generation probability as a function of lllCDR3 connections in individuals (Pn). The contour plots represent lllCDR3s from TCRs identified as virus-specific based on tetramer identification14,15 (red) or with unknown specificity (grey). The Pearson correlation coefficient “R” and p-value “p” are calculated for each group. The black dotted lines delimit the threshold for the 2.5% sequences with both higher Pgen and degree of connection. lllCDR3s with both the highest Pgen and connections were also the most virus-specific for 11 out of 12 individuals (p-value <0.0001, two-tailed Chi-square test). (B) Virus-specific CDR3s among DPCD3^+^ and ThyCD8 cells. Barplots represent the percentage of CDR3s from TCR identified as virus-specific based on tetramer binding(*27, 28*). For each panel, calculation is done on dispersed (left boxplot) or clustered (right boxplot) CDR3s. Colours correspond to different viral specificities.

**Figure S6.**
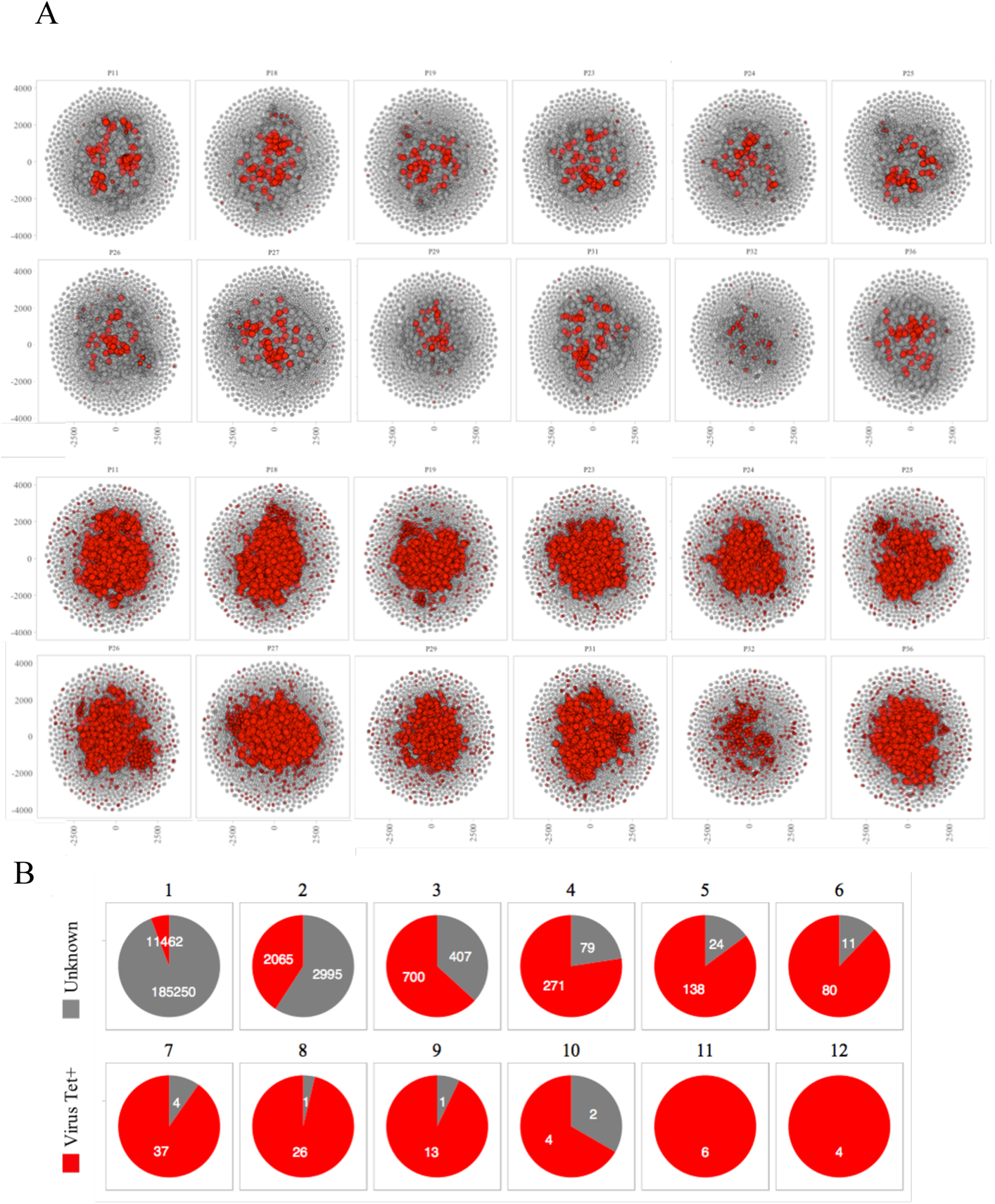
Sharing of virus specific thymocytes’ CDR3s. (A) Network of clustered nodes from the first top ranking 18,000 CDR3s. For each individual, we plot CDR3s as viral-associated (red) either if they are in the public database (line 1 and 2) or if they have a LD≤ 1 with a CDR3 of the public database (line 3 and 4). (B) Sharing of virus-associated CDR3s. The virus-associated CDR3s are highly shared between individuals.

**Figure S7.**
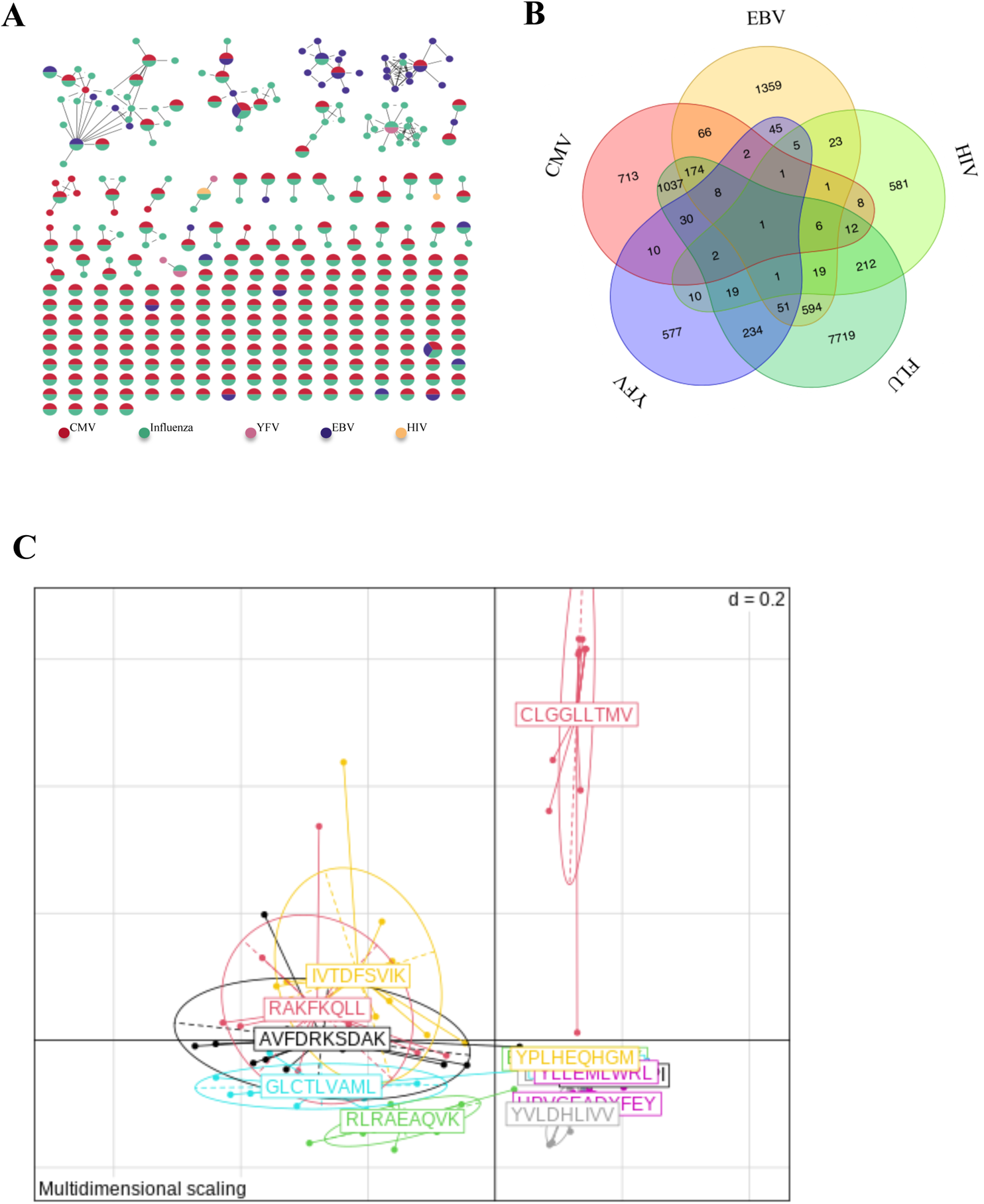
Binding properties of single cell TCRs. A. Network of tetramer-binding CDR3s from public database (VDJdb) identifying CDR3s with multiple viral specificities. Each dot represents a single CDR3. Dot are connected by edges defined by Levenshtein distance of ≤1 (one AA substitution/insertion/deletion). Each dot represents a CDR3 with at least 2 specificities for 2 different viruses. The colors are related to a specific virus. The specificities of these CDR3 were identified by tetramer staining.
B. Venn diagram representing specificities for 5 different viruses of the 13,557 unique virus-associated CDR3s (those from Figure 3e, LD≤1).
C. MDS representation of MH dissimilarity index for EBV repertoire. The MH index are calculated for each repertoire specific of EBV antigen (n=16). Each point represents a patient and each color an antigen.

**Figure S8.**
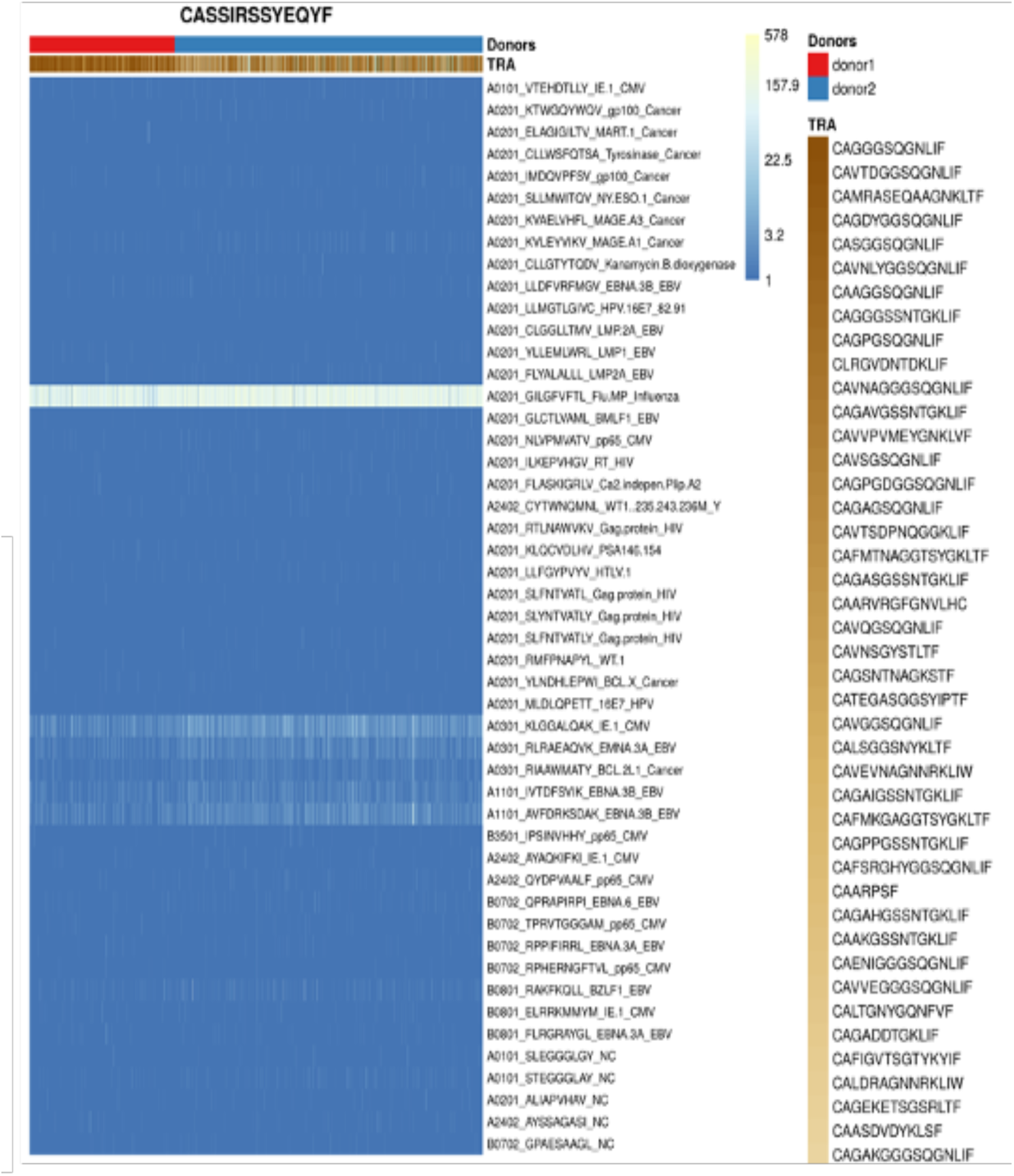
βCDR3s drive polyspecificity. Heatmap of the binding score for different dextramers of different TCRs using the same CASSIRSSYEQYF βCDR3, in two donors. The recognition properties are remarkably similar whatever the βCDR3. The polyspecificity is oriented towards the recognition of common viruses such as influenza, EBV, CMV.

**Figure S9.**
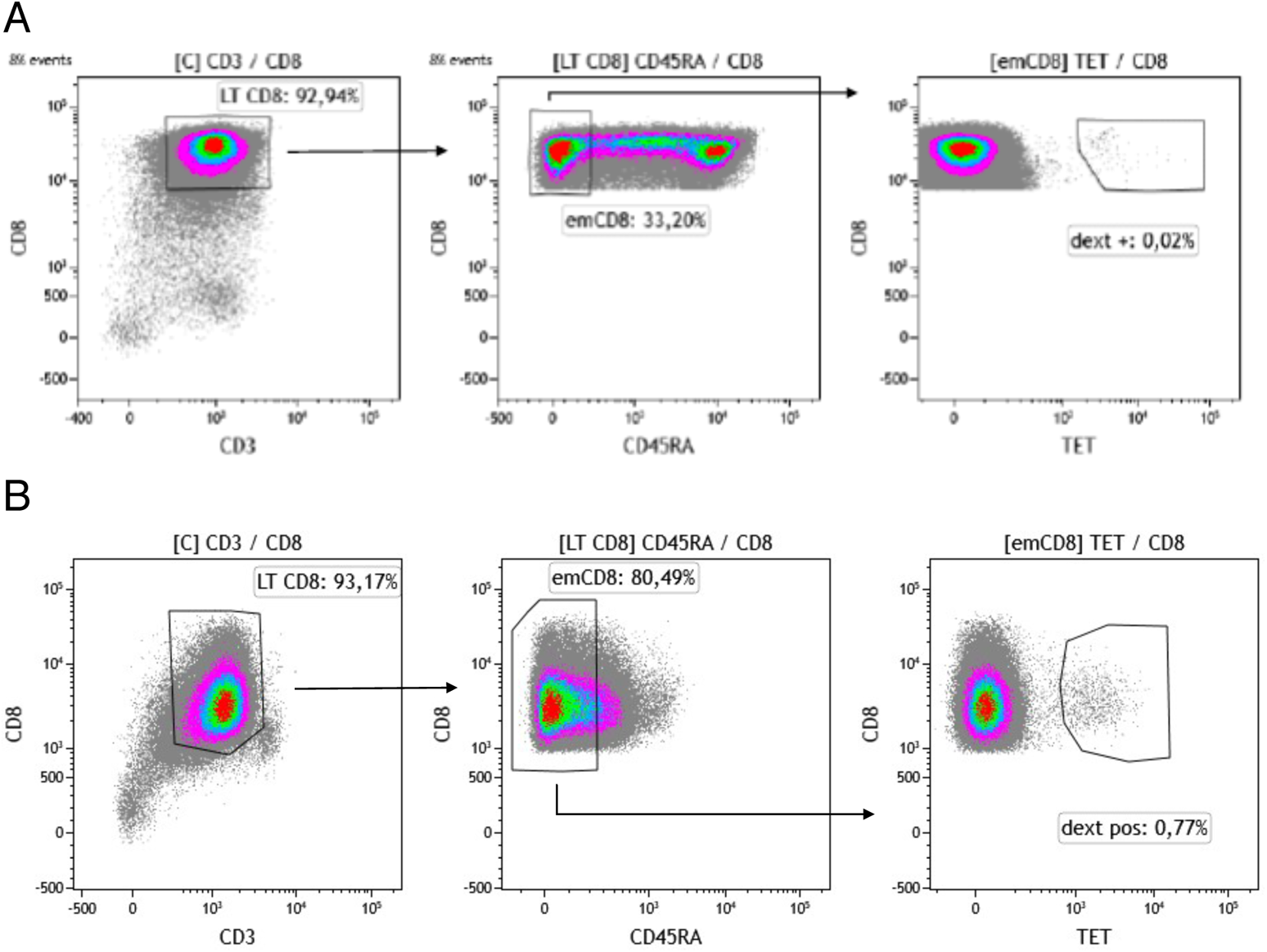
Gating strategy for sorting effector memory (emCD8) Dextramer positive cells. CD8 were previously enriched from cytapheresis PBMC. Gating strategy of the sorting of Cytomegalovirus (A) or Epstein Barr Virus (B) dextramère positive effector memory CD8 (emCD8) from CD8 enriched fraction of PBMC’s donor after excluding doublets. emCD8 cells are CD3^+^ CD8^+^ CD45RA^−^ cells.

**Figure S10.** Polyspecific properties of anti-CMV cytotoxic cells. **(A)** PBMC were stimulated in vitro using Peptivator CMV pp65 (that consists mainly of 15-mer peptides with 11-amino acid overlap, covering the complete sequence of the pp65 protein) and cultured for 25 days. Selected CTL were then restimulated in the presence of either Peptivator CMV pp65, Peptivator CMV IE-1, EBV-BZLF1 or Peptivator EBV EBNA-1. **(B)** Polyspecificity is dependent of peptide level. We represent the % of cells secreting IFNγ as a function of peptides concentration. For pp65 CMV peptide (red) and BZLF-1 EBV peptide (blue), the level of IFN is different and depends of the concentration of the peptides, suggesting different affinity for the peptides. These observations were made with cells from 5 different donors.

**Supplementary table 1.** Enrichment of public βCDR3s in CD8+ thymocytes vs DPCD3+. Contingency table for the Chi-square analysis performed with Yates’ correction to test the null hypothesis of independence between the sharing of βCDR3s (Public or Private) vs the cell phenotype (DPCD3^+^ and CD8^+^). We performed this test in the 10 donors for which we have 18,000 βCDR3s in both DPCD3^+^ and CD8^+^ thymocytes. A βCDR3 is defined as public if it is found at least once in the 18,000 βCDR3s of the same cell phenotype from other donors. The results (p-value < 0.0001) rejected the null hypothesis, thereby indicating the interdependency of the two variables.

|  |  | DP | CD8 | p-value | Odds ratio |
| --- | --- | --- | --- | --- | --- |
| P11 | Public | 590 | 1630 | p<0.0001 | 0.3403 |
|  | Private | 17410 | 16370 |  |  |
| P18 | Public | 695 | 1567 | p<0.0001 | 0.4212 |
|  | Private | 17305 | 16433 |  |  |
| P19 | Public | 485 | 1489 | p<0.0001 | 0.3071 |
|  | Private | 17515 | 16511 |  |  |
| P23 | Public | 649 | 1489 | p<0.0001 | 0.4148 |
|  | Private | 17351 | 16511 |  |  |
| P24 | Public | 578 | 1416 | p<0.0001 | 0.3886 |
|  | Private | 17422 | 16584 |  |  |
| P25 | Public | 640 | 1536 | p<0.0001 | 0.3952 |
|  | Private | 17360 | 16464 |  |  |
| P26 | Public | 584 | 1470 | p<0.0001 | 0.3771 |
|  | Private | 17416 | 16530 |  |  |
| P27 | Public | 640 | 1583 | p<0.0001 | 0.3823 |
|  | Private | 17360 | 16417 |  |  |
| P29 | Public | 655 | 1583 | p<0.0001 | 0.3916 |
|  | Private | 17345 | 16417 |  |  |
| P36 | Public | 653 | 1468 | p<0.0001 | 0.4239 |
|  | Private | 17347 | 16532 |  |  |

**Supplementary table 2.** Enrichment of virus-specific lllCDR3s from databases14,15 in clustered CD8+ thymocytes. Contingency table for the Chi-square analysis performed with Yates’ correction to test the null hypothesis of independence between the specificity of βCDR3s (Virus Tet+ and Unknown specificities) vs the connection of βCDR3 (“clustered” and “dispersed”) in all the CD8+ thymocytes from 12 donors. The results (p-value < 0.0001) rejected the null hypothesis, thereby indicating the interdependency of the two variables.

|  |  | Clustered | Dispersed | p-value | Odds ratio |
| --- | --- | --- | --- | --- | --- |
| P11 | Virus Tet+ | 106 | 8 | p<0.0001 | 40.75 |
|  | Unknown | 4389 | 13497 |  |  |
| P18 | Virus Tet+ | 126 | 10 | p<0.0001 | 39.93 |
|  | Unknown | 4285 | 13579 |  |  |
| P19 | Virus Tet+ | 102 | 8 | p<0.0001 | 46.77 |
|  | Unknown | 3832 | 14058 |  |  |
| P23 | Virus Tet+ | 93 | 7 | p<0.0001 | 47.82 |
|  | Unknown | 3892 | 14008 |  |  |
| P24 | Virus Tet+ | 96 | 9 | p<0.0001 | 40.38 |
|  | Unknown | 3739 | 14156 |  |  |
| P25 | Virus Tet+ | 112 | 7 | p<0.0001 | 63.60 |
|  | Unknown | 3594 | 14287 |  |  |
| P26 | Virus Tet+ | 83 | 2 | p<0.0001 | 146.1 |
|  | Unknown | 3963 | 13952 |  |  |
| P27 | Virus Tet+ | 99 | 8 | p<0.0001 | 34.60 |
|  | Unknown | 4714 | 13179 |  |  |
| P29 | Virus Tet+ | 55 | 9 | p<0.0001 | 31.65 |
|  | Unknown | 2903 | 15033 |  |  |
| P31 | Virus Tet+ | 104 | 1 | p<0.0001 | 360.8 |
|  | Unknown | 4004 | 13891 |  |  |
| P32 | Virus Tet+ | 44 | 9 | p<0.0001 | 32.93 |
|  | Unknown | 2320 | 15627 |  |  |
| P36 | Virus Tet+ | 86 | 6 | p<0.0001 | 54.12 |
|  | Unknown | 3750 | 14158 |  |  |

**Table S3.**
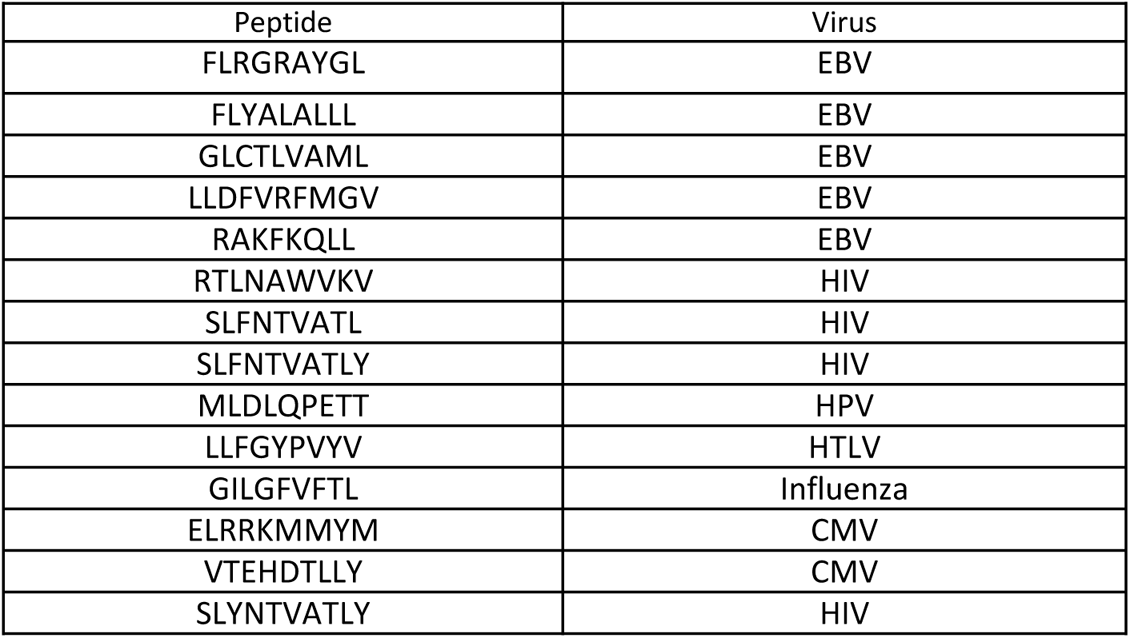
List of peptides represented on the chord plot. Related to 4G right. The table is organized according to the clockwise order of the chord plot segments.

